# White matter microstructure links with brain, bodily and genetic attributes in adolescence, mid- and late life

**DOI:** 10.1101/2024.11.27.625603

**Authors:** Max Korbmacher, Mario Tranfa, Giuseppe Pontillo, Dennis van der Meer, Meng-Yun Wang, Ole A. Andreassen, Lars T. Westlye, Ivan I. Maximov

## Abstract

Advanced diffusion magnetic resonance imaging (dMRI) allows one to probe and assess brain white matter (WM) organization and microstructure *in vivo*. Various dMRI models with different theoretical and practical assumptions have been developed, representing partly overlapping characteristics of the underlying brain biology with potentially complementary value in the cognitive and clinical neurosciences. To which degree the different dMRI metrics relate to clinically relevant geno-and phenotypes is still debated. Hence, we investigate how tract-based and whole WM skeleton parameters from different single- and multi-compartment dMRI approaches associate with clinically relevant and white matter-related phenotypes (sex, age, pulse pressure (PP), body-mass-index (BMI), brain asymmetry) and genetic markers in the UK Biobank (UKB, n=52, 140) and the Adolescent Brain Cognitive Development (ABCD) Study (n=5, 844). Multi-compartment dMRI approaches provided the strongest WM associations with age, and unique insights into brain asymmetry. Kurtosis was most indicative of PP and BMI. WM-based sex classifications and polygenic score associations for common psychiatric disorders and Alzheimer’s disease were similar across diffusion approaches. We conclude that WM microstructure is differentially associated with clinically relevant pheno- and genotypes at different points in life. Multi-compartment dMRI approaches, and particularly the examined Bayesian approach, provide additional information to conventional approaches in such examinations.

## 1 Introduction

White matter (WM) is characterised by bundles of axonal neurites that stretch across the human brain. By connecting grey matter neurons with each other, fibre bundles allow for neural communication across the brain. Diffusion magnetic resonance imaging (dMRI) probes *in vivo* brain organization and microstructure, and is a particularly powerful tool for WM mapping. DMRI allows for quantification and visualisation of different brain tissue features. Recent advances (Fieremans et al., 2011; Kaden, Kelm, et al., 2016; Kaden, Kruggel, & Alexander, 2016; Novikov, Veraart, et al., 2018; Novikov et al., 2019; Reisert et al., 2017) have led to many proposed approaches to model and characterise WM beyond commonly used diffusion tensor imaging (DTI) (Novikov, Kiselev, & Jespersen, 2018) and approaches exploiting non-Gaussian diffusivity (kurtosis) (Fieremans et al., 2011; Jensen et al., 2005). However, such approaches are limited by not being able to address the complexity of the tissue structure, for instance, crossing fibres or microscopic cellular features (Dell’Acqua & Tournier, 2019; Novikov et al., 2019). Hence, new approaches were developed to more adequately assess WM microstructure. These approaches generally rely on the standard model of diffusion, which separates intra- and extra-axonal space (Novikov, Veraart, et al., 2018; Novikov et al., 2019; Reisert et al., 2017), with different assumptions about the characteristics of these intra- and extra-axonal compartments (e.g., modeling diffusion within impermeable cylinders or spheres of different sizes). Based on such assumptions, one can derive a range of scalar parameters reflecting different WM features within the framework of the standard diffusion model or empiric diffusion representation. As a result, there is a multitude of diffusion parameters often having closely related meaning or presenting strong correlations between approaches (Beck et al., 2021; Korbmacher, de Lange, et al., 2023). For each parameter, one can create voxel-wise maps or average across WM tracts and regions. Thus, these scalar maps offer quantitative features of WM organization and structure.

As the parameters across dMRI approaches describe diffusion, they are also related to each other. However, their sensitivity to and complementary value for predicting clinically relevant traits is unclear. Mapping such relationships can inform future selections of dMRI approaches to extract meaningful metrics to answer specific research questions about the role of WM in health and disease.

Hence, in the present comparative analyses, we present effect sizes (associations and group differences) for a range of clinically relevant metrics and the parameters of six dMRI approaches in two large-scale MRI datasets, the UK Biobank (UKB) and the Adolescent Brain Cognitive Development (ABCD) Study. The non-WM metrics were chosen based on previous evidence indicating WM-associations, and their general clinical relevance for cardiometabolic, neurological and psychiatric disorders. These metrics include sex, age, brain asymmetry, body-mass-index, pulse pressure, and polygenic risk scores. We highlight the strongest effects to assess dMRI approaches as a whole (such as Diffusion Tensor Imaging), spatial specificity on the tract-level (such as the cingulum-hippocampus tract), and metric-specificity (e.g., the intra-axonal water fraction from the Bayesian multi-compartment approach).

## 2 Methods

### 2.1 Samples

We analysed diffusion data from two big MRI studies: UK Biobank (Miller et al., 2016) and ABCD (Casey et al., 2018). We obtained *N*_*total*_ = 52, 140 datasets, with *N*_*ABCD*_ = 5, 844 and *N*_*UKB*_ = 46, 196. Due to variability in ethnicity, relevant to polygenic assessments, we controlled for participants’ ethnicity in all genotype-related analyses.

### 2.2 Diffusion data

We processed 26 parameters (see for overview Appendix M) from the following diffusion approaches: the Bayesian Rotationally Invariant Approach (BRIA) (Reisert et al., 2017), Diffusion Kurtosis Imaging (DKI) (Jensen et al., 2005), Diffusion Tensor Imaging (DTI) (Basser et al., 1994), the Spherical Mean Technique (SMT) (Kaden, Kruggel, & Alexander, 2016), and its multicompartment extension (SMTmc) (Kaden, Kelm, et al., 2016), and White Matter Tract Integrity (WMTI) (Fieremans et al., 2011).

Quality control comprised of the YTTRIUM method (Maximov et al., 2021) which converts global dMRI scalar metrics into 2D format using a structural similarity extension of each scalar map to their mean image in order to create a 2D distribution of image and diffusion parameters. Non-clusterised values are then excluded. Additional exclusions entailed tract-based values exceeding 5 standard deviations from the mean in each parameter. This procedure led to 697 exclusions in ABCD (11.93%) and 2,555 in the UKB data (5.53%). We obtained a final sample of *N*_*total*_ = 42, 230, with *N*_*ABCD*_ = 5, 147 and *N*_*UKB*_ = 37, 083.

### 2.3 MRI acquisition and post-processing

UKB MRI data acquisition procedures and protocols are described elsewhere (Alfaro-Almagro et al., 2018; Miller et al., 2016; Sudlow et al., 2015). For ABCD acquisition protocols see org/scientists/protocols/. The dMRI protocols in these studies are different and optimised for different diffusion approaches.

After access to the raw dMRI data, we processed the data using an optimised pipeline (Maximov et al., 2019), including corrections for noise (Veraart et al., 2016), Gibbs ringing (Kellner et al., 2016), susceptibility-induced and motion distortions, and eddy current artifacts (Andersson & Sotiropoulos, 2016). Isotropic 1 mm^3^ Gaussian smoothing for UKB and 0.8 mm^3^ in ABCD was carried out using FSL’s (Jenkinson et al., 2012; Smith et al., 2004) *fslmaths* (FSL version 6.0.1). Employing the multi-shell data, DTI, DKI (Jensen et al., 2005) and WMTI (Fieremans et al., 2011) parameters were estimated using Matlab 2017b code (https://gitgub.com/NYU-DiffusionMRI/DESIGNER). SMT (Kaden, Kruggel, & Alexander, 2016), mcSMT (Kaden, Kelm, et al., 2016) parameters were estimated using original (Kaden, Kelm, et al., 2016; Kaden, Kruggel, & Alexander, 2016) in-house C++ code (https://github.com/ekaden/smt). BRIA estimates were evaluated with the original (Reisert et al., 2017) Matlab code (https://bitbucket.org/reisert/baydiff/src/master/).

We used Tract-based Spatial Statistics (TBSS) (Smith et al., 2006), as part of FSL (Jenkinson et al., 2012; Smith et al., 2004). Initially, all brain-extracted (Smith, 2002) fractional anisotropy (FA) images were aligned to MNI space using non-linear transformation (FNIRT) (Jenkinson et al., 2012). Next, the mean FA image and mean FA skeleton were derived. Each diffusion scalar map was projected onto the mean FA skeleton using TBSS. To provide a quantitative description of diffusion parameters at the tract level, we used the John Hopkins University (JHU) atlas (Hua et al., 2008; Mori et al., 2006) 20 tract averages based on a probabilistic WM atlas for each of the 26 parameters, totalling 520 values per individual. We additionally computed the skeleton average for each of the 26 parameters.

### Polygenic risk scores (PGRS)

We estimated PGRS for each participant with available genomic data, using LDPred2 (Privé et al., 2020) with default settings. As input for the PGRS, we used summary statistics from recent genome-wide association studies of Autism Spectrum Disorder (ASD) (Autism Spectrum Disorders Working Group of The Psychiatric Genomics Consortium, 2017), Major Depressive Disorder (MDD) (Wray et al., 2018), Schizophrenia (SCZ) (Trubetskoy et al., 2022), Attention Deficit Hyperactivity Disorder (ADHD) (Demontis et al., 2019), Bipolar Disorder (BIP) (Mullins et al., 2021), Obsessive Compulsive Disorder (OCD) (Arnold et al., 2018), Anxiety Disorder (ANX) (Otowa et al., 2016), and Alzheimer’s Disease (AD) (Wightman et al., 2021). We used a minor allele frequency of 0.05, as the threshold most commonly used in PGRS studies of psychiatric disorders.

### 2.4 Cardiometabolic risk factors

We used a selection of cardiometabolic factors, which have previousy been linked to WM (Beck et al., 2022; Korbmacher, Gurholt, et al., 2023; Korbmacher, van der Meer, Beck, de Lange, et al., 2024), and which were available in both datasets. Body-mass-index (BMI) was calculated as *weight*(*kg*)*/*(*height*(*m*))^2^ and PP (mmHg) as the difference between systolic and diastolic PP. For a better estimation of PP, we used the average of PPs derived from each automated reading.

### 2.5 Statistical analyses

First, we correlated all skeleton and tract-level average metrics with each other for a descriptive overview of associations between metrics across dMRI approaches.

Second, we used the WM parameters from each dMRI approach to classify participants’ into males or females based on their tract-level metrics (*M*_1_ to *M*_*i*_) using logistic regression. Due to multicollinearity among the predictors, we used principal components analyses to obtain orthogonal predictors. As the 5 first components (*PC*_1_ to *PC*_5_) of the tract-level scalars of each dMRI approach explained between 60% and 90% of the variance, we limited the analysis to 5 PCs for these classifications and further analyses. This was leading to 7 classifications per dataset and additionally 7 classifications for the combined data (14 classifications in total).

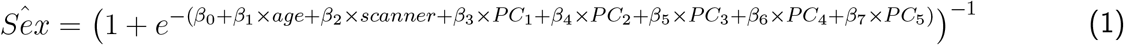

To add explainability to the principal components (used across analyses), we correlated each of the resulting five components (*PC*_*j*_) with each tract-level metric (*M*_*i*_) individually while controlling for covariates (scanner, sex, age) using simple linear regression models (*N*_*tests*_ = 5(*PCs*) × 26(*metrics*) = 130). This approach was chosen in order to being able to correct for covariates:

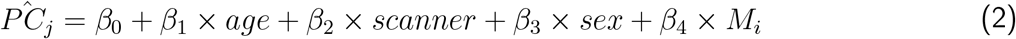

Third, we associated the aforementioned principal components (*P C*_*j*_) and age, as well as skeleton-averaged metrics (*S*_*l*_, *N*_*associations*_ = 7(*PC*) + 26(*S*) = 33):

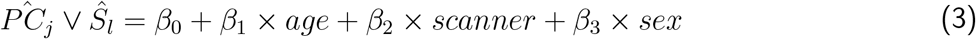

Fourth, we assessed tract asymmetries using paired-sample t-tests on each metric for each pair of lateralised tracts (*N*_*t*−*tests*_ = 9(*tracts*) × 26(*metrics*) = 234).

Fifth, we assessed associations between the cardiovascular risk factors BMI and PP (PP) and the mentioned principal components (*PC*_*j*_) and skeleton averages (*S*_*l*_,*N*_*associations*_ = 2(*BMI* V *PP*) ×(7(*PC*)+ 26(*S*)) = 66):

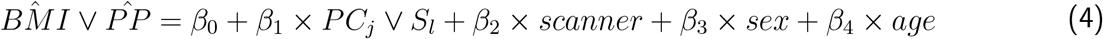

Sixth, we assessed associations between PGRS and principal components (*PC*) and skeleton averages (*S*). For that, we initially estimated the first principal component of the PGRS for common psychiatric disorders. For the associations between the PGRS for Alzheimer’s disease, we used the raw value (*N*_*associations*_= 7 (*PGRS*)× 33 (7 tract level models + 26 skeleton averages) = 231). To avoid con-founding genetic effects of ethnicity, we also included ethnicity as a factor:

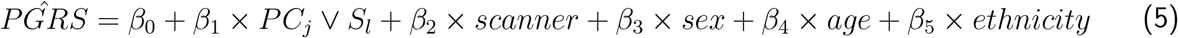

All analyses were executed on a) the UKB data, b) the ABCD data. There was only one exception where both datasets were analysed together: the correlations between the diffusion parameters, to showcase that the associations were robust, independent of the data composition. The data were scaled by subtracting the mean and dividing by the standard deviation, and standardised *β* coefficients were reported.

A total number of 708 analyses per dataset were conducted. We adjusted our *α*-level accordingly, using a conservative Bonferroni correction to control false positives, leading to *α* = 0.05*/*708 = 0.000071.

Considering the tests with lowest power defined by 4 numerator degrees of freedom (see formulas above) and 4,625 denominator degrees of freedom (based on the smallest sample, the ABCD data, testing for BMI and PP), at an adjusted significance level of 0.0071% and 95% power, one can detect effects as low as Cohen’s *f* ^2^ = 0.0086, which closely translates to a coefficient of determination of *R*^2^ = 0.0086 or a correlation coefficient of *r* = 0.093, and a Cohen’s *d* = 0.186. Using the same parameters yielded minimum detectable effect sizes in the UK Biobank of *f* ^2^ ≈ *R*^2^ = 0.00125, a correlation coefficient of *r* = 0.0354, and Cohen’s *d* = 0.071. All analyses were conducted in R, version 4.2.1.

## 3 Results

Skeleton-averaged metrics correlated conditionally in two clusters, describing a) radial and axial diffu-sivity b) FA and kurtosis. Notably, dependencies of water fraction metrics from the different diffusion approaches varied in their relationships with other metrics dependent on different diffusion protocols. Hence, partial voluming cannot be excluded in the presence of the free water compartment (Fig. 1).

**Figure 1.**
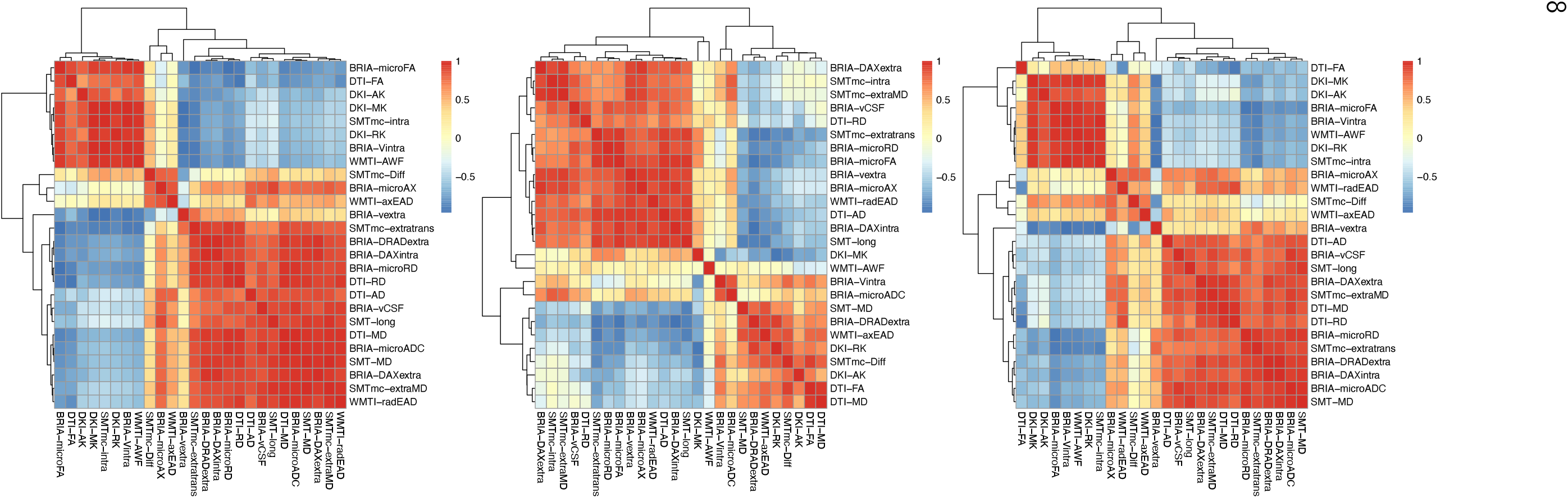
Correlations between skeleton-level averaged metrics from each of the diffusion approaches. Panels from left to right indicate UKB, ABCD, and merged data (UKB and ABCD data). Correlations were clustered both column- and row-wise. BRIA = Bayesian Rotationally Invariant Approach. DKI = Diffusion Kurtosis Imaging. DTI = Diffusion Tensor Imaging. SMT = Spherical Mean Technique. SMTmc = SMT’s multi-compartment version. WMTI = WM Tract Integrity. Metrics for each of the approaches are specified in Appendix

Due to multicollinearity, we used the first five principal components of the respective dMRI approaches’ tract-level estimates, instead of raw tract scores. The first two principal components generally explained together more than 60% of the variance, with the exception of WMTI in the ABCD data, where the first three components explained more than 60% of the variance of the individual tract-level scalars. Scree plots showing the variance explained for each of the approaches and their combination can be found in Appendices A-B.

### 3.1 Sex classifications

The accuracy for sex classifications from tract-level scalars was similar across approaches, yet with DKI and BRIA performing slightly better in ABCD data and DTI and WMTI in UKB data (Fig. 2). The contribution of the principal components to the predictions was strongest represented in the fourth and fifth components, which generally explained less of the variance across tract metrics than the first components (see Appendices A-B). The corrected associations of tract-level metrics with the fourth and fifth components of the different approaches highlighted a spatially distributed pattern, yet with higher loadings assigned to cingulate gyrus and cingulate-hippocampus tracts in UKB PC4 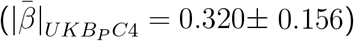 and ABCD PC5 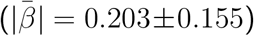, and uncinate fasciculus for UKB PC5 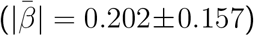 and ABCD PC4 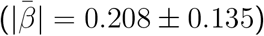 relative to the other loadings (see Appendices and K-L).

**Figure 2.**
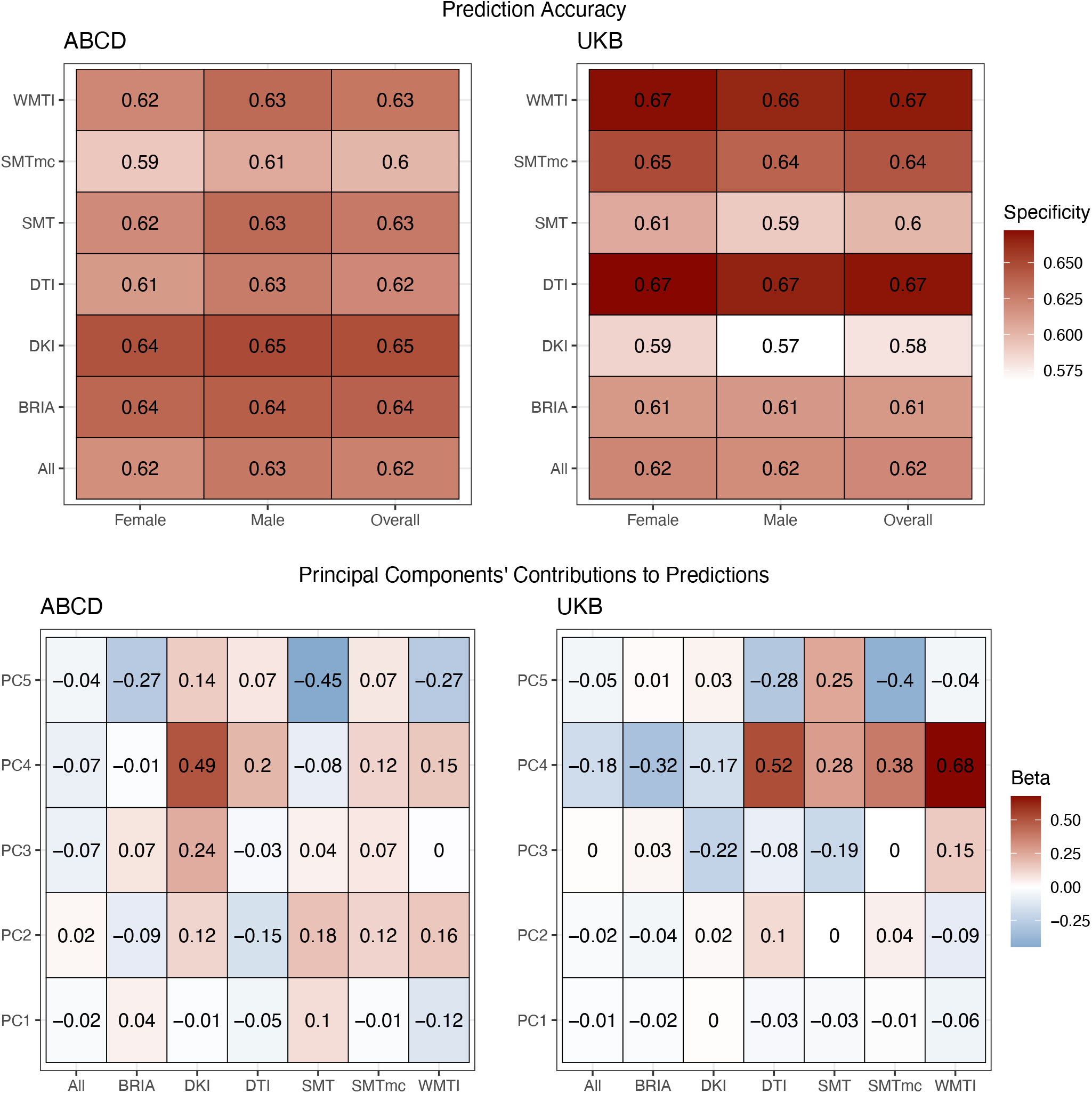
Sex prediction accuracy and respective principal components’ contributions to the predictions. The top row panels indicate the prediction accuracy. The bottom two panels indicate the contribution of each of the five utilised principal components to the sex classification from each of the dMRI approaches indicated by age and scanner corrected correlations. Standardised beta coefficients are presented. All models were significant at *p*_*adj*_ *<* .001. DKI = Diffusion Kurtosis Imaging. DTI = Diffusion Tensor Imaging. SMT = Spherical Mean Technique. SMTmc = SMT’s multi-compartment version. WMTI = White Matter Tract Integrity.

### 3.2 Age associations

Principal components of BRIA tract-level metrics explained most of the variance in age (ABCD: *R*^2^ = 12%, UKB: *R*^2^ = 36%). While DKI and SMTmc were explaining the same share of age variance as BRIA in the adolescence sample (*R*^2^ = 12%), DTI and WMTI explained more variance in the mid-to late life sample (DTI: *R*^2^ = 33%, WMTI: *R*^2^ = 30%; Fig. 3). The third to fifth principal components, showing loadings across metrics and tracts, were most relevant for age predictions across approaches. The third and fourth components were the only components to present sufficiently powered age-associations (*r*_*min*_ = 0.09) for SMT in the ABCD sample, with SMT-long loading strongest in forceps major (*β* = 0.41 ± 0.004) and cingulate gyrus (*β >* 0.355) in the UKB and anterior thalamic radiation in ABCD (*β >* 0.431) on the third component and the corticospinal tract on the fourth component (*β >* 0.285) in UKB and cingulate gyrus (*β >* 0.378) and forceps minor (*β* = 0.387 ± 0.013) in ABCD. For the larger and older sample (UKB), multiple age-associations were sufficiently powered (*r*_*min*_ = 0.04). The most relevant component (four) in the best performing model (BRIA) loaded strongest on the cingulate gyrus tract for intra- and extra-axonal radial diffusivities (*β >* 0.408) in UKB, and forceps minor cerebrospinal fluid fraction (*β* = −0.460 ± 0.010), microscopic ADC (*β* = −0.388 ± 0.009), and microscopic radial diffusivity in ABCD (*β* = −0.356 ± 0.008; Appendices C-L). Direct associations between skeleton-level scalars and age present mostly opposing trends when comparing adolescence to the mid-to late life samples (ABCD and UKB). BRIA’s cerebrospinal fluid fraction and microscopic metrics which mimic DTI metrics on the microstructure level presented the strongest age associations (Fig. 4). At the same time, BRIA’s intra and extra-axonal water fractions were among the largest age-associations during adolescence, together with kurtosis metrics.

**Figure 3.**
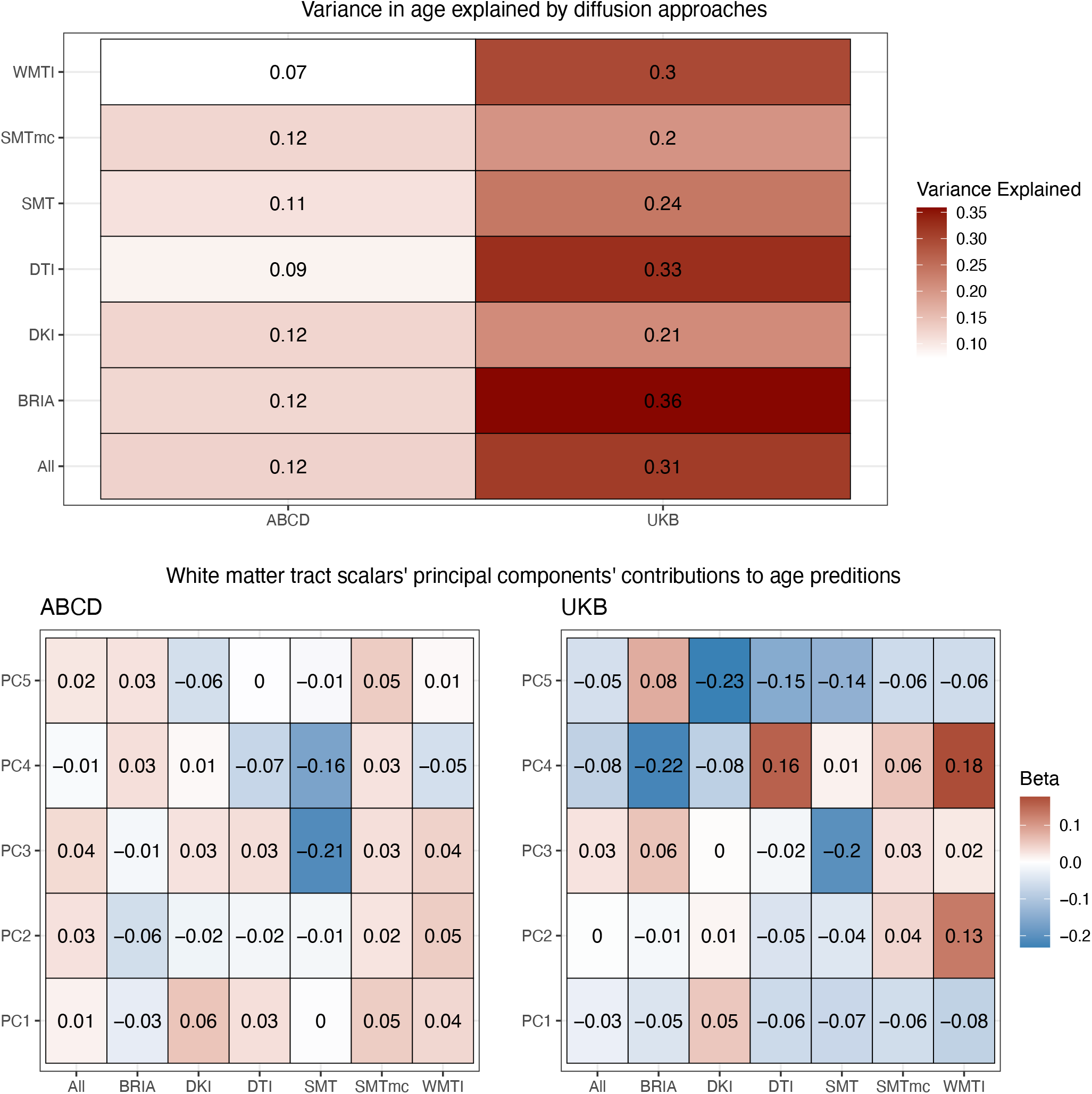
Variance in age explained by principal components of the different diffusion approaches (top) and each component’s contribution to age predictions indicated by age and scanner corrected correlations (bottom). Standardised beta coefficients are presented. DKI = Diffusion Kurtosis Imaging. DTI = Diffusion Tensor Imaging. SMT = Spherical Mean Technique. SMTmc = SMT’s multi-compartment version. WMTI = White Matter Tract Integrity.

**Figure 4.**
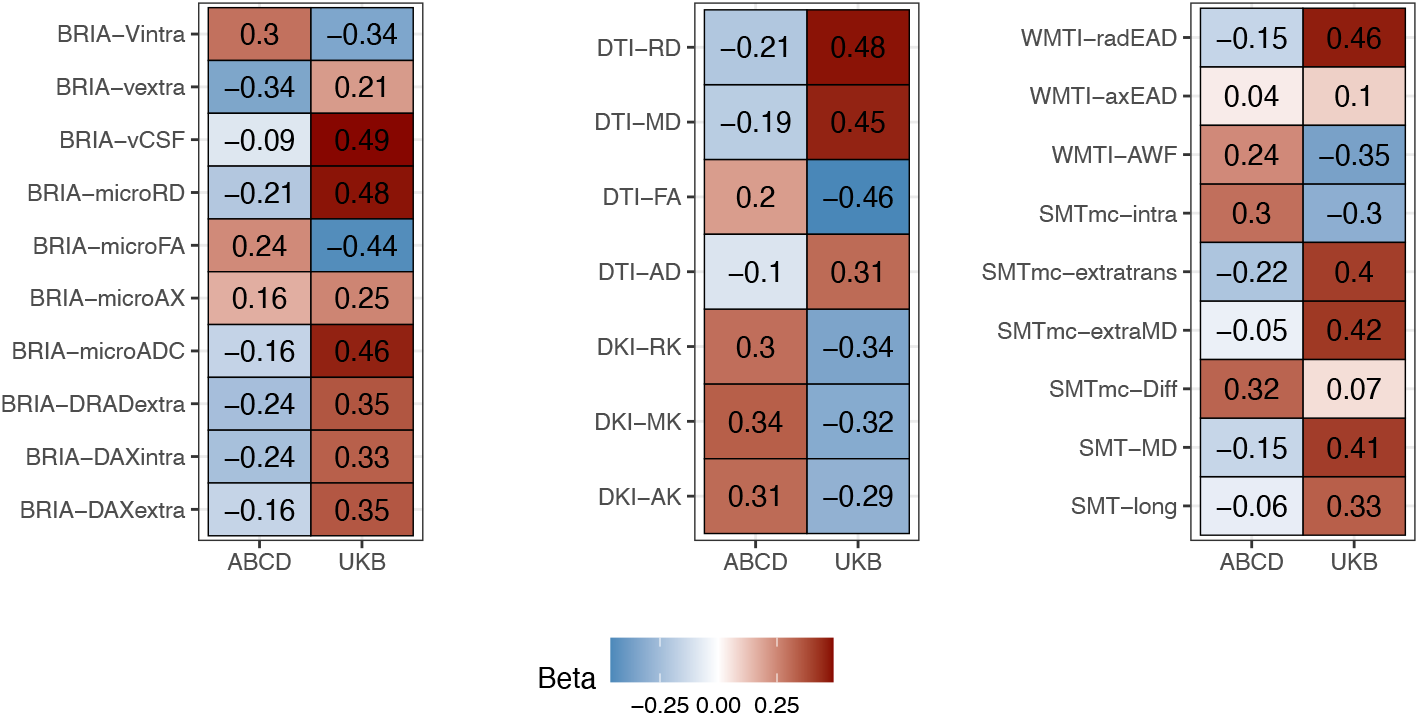
Skeleton averaged scalars’ age associations in ABCD and UKB data. Standardised beta coefficients are presented. BRIA = Bayesin Rotationally Invariant Approach. DKI = Diffusion Kurtosis Imaging. DTI = Diffusion Tensor Imaging. SMT = Spherical Mean Technique. SMTmc = SMT’s multi-compartment version. WMTI = White Matter Tract Integrity. Metrics for each of the approaches are specified in Appendix M. Only age associations for BRIA vCSF in the ABCD data set were not sufficiently powered (r *<* 0.09) and all other associations were significant (p *<* 0.001).

### 3.3 Tract asymmetries

We show that the left hemisphere generally presents larger effect sizes, i.e., leftward asymmetry or left lateralisation (Fig. 5). This leftwards asymmetry is stronger expressed in the UKB data 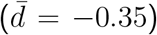 compared to the ABCD dataset 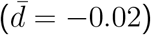 when averaging across effect sizes in each dataset. However, more extreme asymmetries were observed in the ABCD data by DTI and multi compartment approaches BRIA, WMTI and SMTmc. The strongest differences between UKB and ABCD data were captured by BRIA metrics, showing stronger leftward asymmetry in the UKB compared to ABCD dataset (Fig. 5).

**Figure 5.**
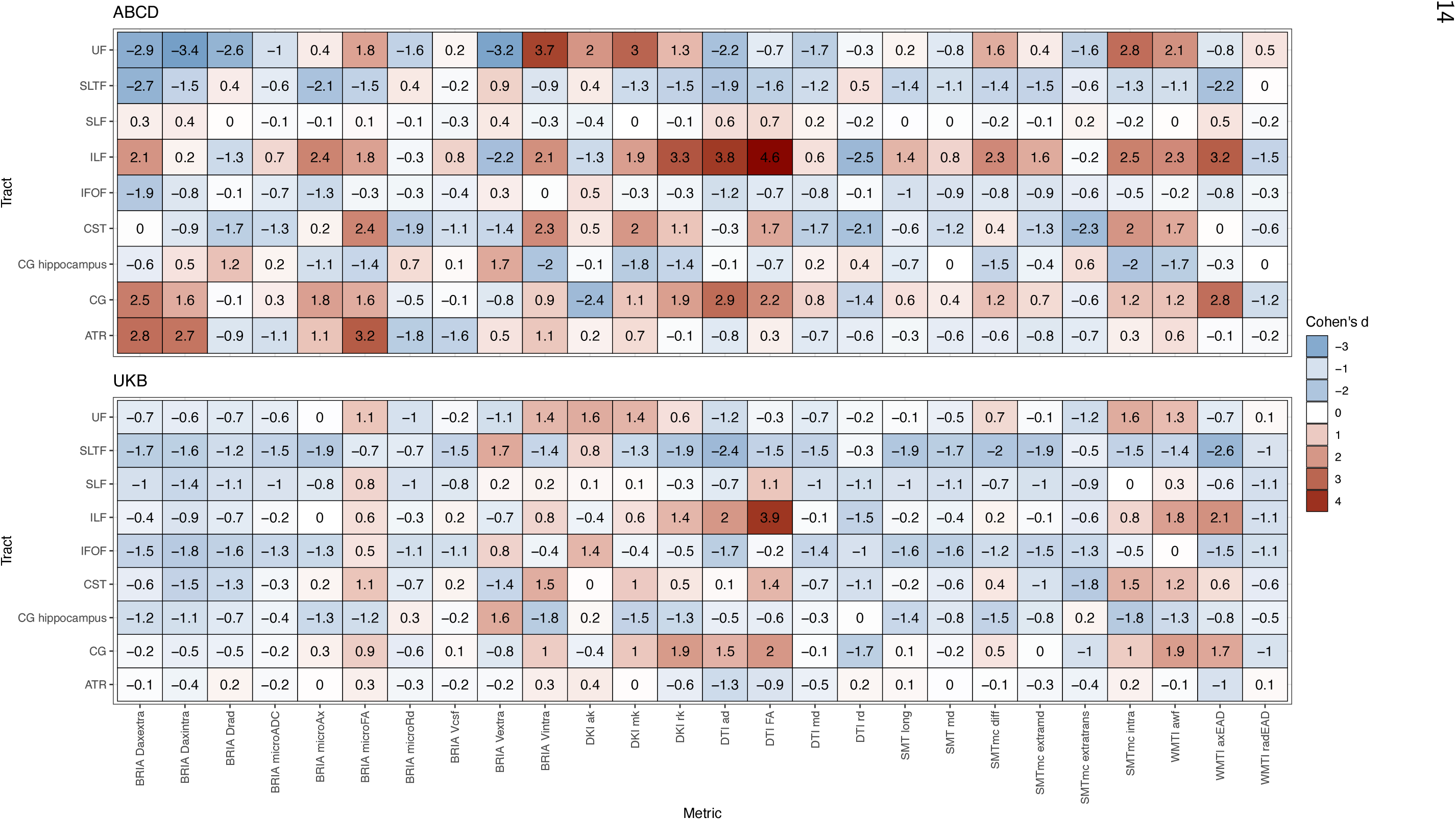
Left-right tract-level differences in ABCD and UKB data. Cohen’s d values are presented. BRIA = Bayesin Rotationally Invariant Approach. DKI = Diffusion Kurtosis Imaging. DTI = Diffusion Tensor Imaging. SMT = Spherical Mean Technique. SMTmc = SMT’s multi-compartment version. WMTI = White Matter Tract Integrity. Metrics for each of the approaches are specified in Appendix M. UF = Uncinate Fasciculus. SLTF = Superior Longitudinal Fasciculus. SLF = Superior Longitudinal Fasciculus (temporal part). ILF = Inferior Lngitudinal Fasciculus. IFOF = Inerior Fronto-Occipial Fasciculus. CST = Cortico-Spinal Tract. CG hippocampus = Cingulum (hippocampus). CG = Cingulum (cinglate gyrus). ATR = Anterior Thalamic Radiation. Note: all |*d*|_*ABCD*_ *>* 0.186 and |*d*|_*UKB*_ *>* 0.071 were sufficiently powered.

### 3.4 Body mass index (BMI) and pulse pressure (PP)

As a reference, we used independent samples t-test to compare BMI and PP in the current sample (*N*_*ABCD*_ = 4, 630, *N*_*UKB*_ = 37, 082) to the rest of the ABCD (*N*_*ABCD*_ = 5, 410) and UKB population (*N*_*UKB*_ = 465, 330). The current sample presented lower PP (*d*_*ABCD*_ = −0.115, 95% *CI*[−0.155, −0.075], *p*_*adj*_ = 1.284 × 10^−7^, *d*_*UKB*_ = −0.083, 95% *CI*[−0.104, −0.062], *p*_*adj*_ = 4.613 × 10^−7^) and BMI (*d*_*ABCD*_ = −0.092, 95% *CI*[−0.131, −0.052], *p*_*adj*_ = 3.018 × 10^−5^, *d*_*UKB*_ = −0.236, 95% *CI*[−0.255, −0.217], *p*_*adj*_ = 1.640 × 10^−113^) compared to the rest of the respective samples.

The axial kurtosis (DKI) and diffusivity (DTI) were among the strongest predictors of BMI (Fig. 6). Yet, several microstructure metrics from BRIA and WMTI predicted BMI in ABCD. Skeleton-level averaged metrics could, however, not predict PP in ABCD (smallest observable effect: *r* = 0.093) and were similarly predicted across diffusion approaches, with the exception of DKI, which showed small associations with PP (|*r*| *<* 0.05).

**Figure 6.**
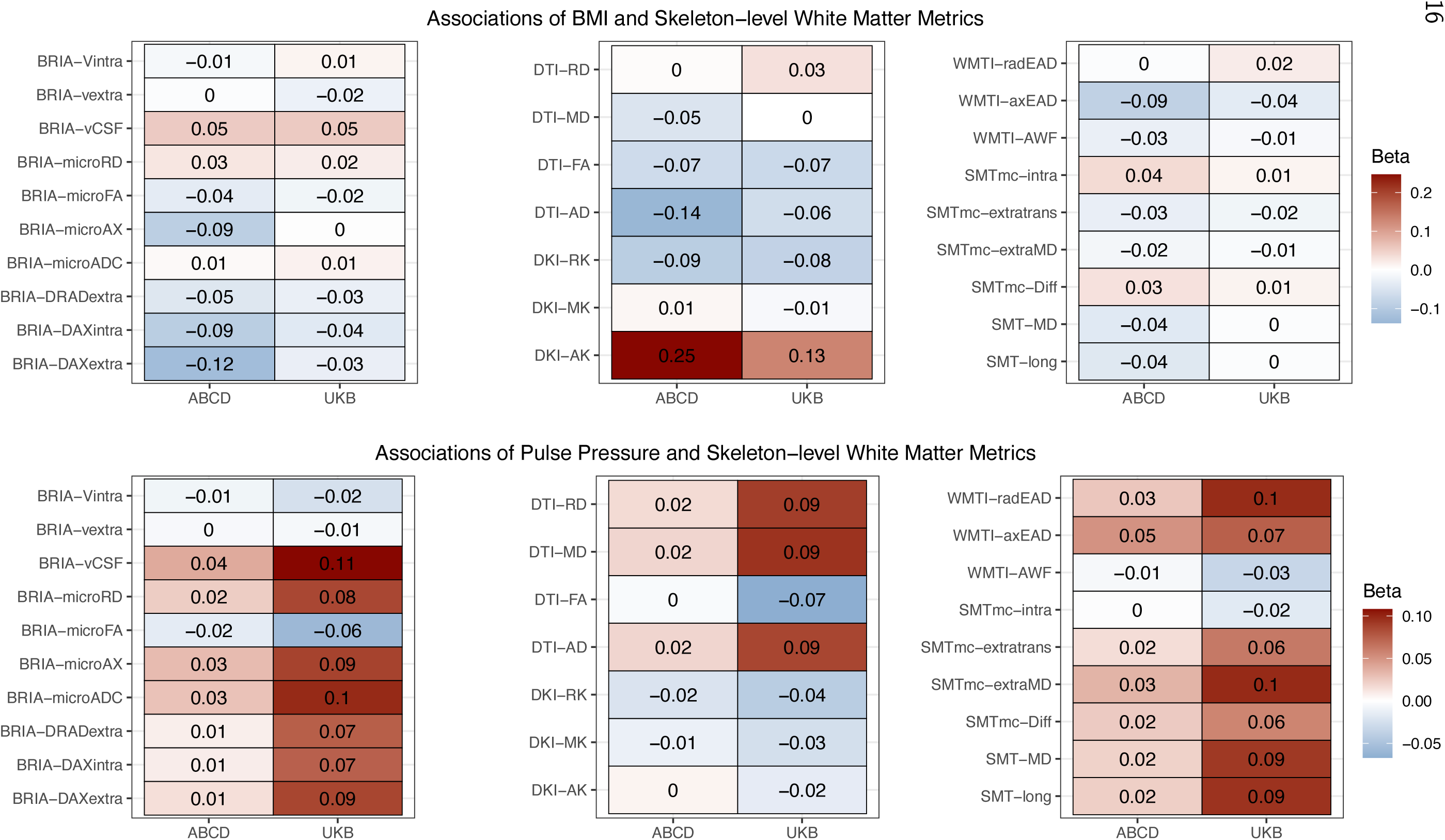
Skeleton-level white matter associations with BMI and PP. Associations are indicated by standardized beta coefficients. BRIA = Bayesin Rotationally Invariant Approach. DKI = Diffusion Kurtosis Imaging. DTI = Diffusion Tensor Imaging. SMT = Spherical Mean Technique. SMTmc = SMT’s multi-compartment version. WMTI = White Matter Tract Integrity. Metrics for each of the approaches are specified in Appendix M. Note: only BRIA, DKI, DTI, and WMTI metrics were sufficiently powered predictors of BMI. PP associations were insufficiently powered in ABCD data.

The skeleton-level associations were reflected when using the first five principal components of the respective dMRI approaches to predict BMI and PP: BMI was similarly predictive across approaches. However, DKI components two and four were adding to the predictions in both UKB and ABCD (Fig. 7). While the fourth component was reflective of the general tract-level variance across metrics, the third component was strongest associated with cingulate, cingulum-hippocampus, and corticospinal tract metrics in UKB data 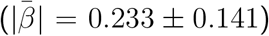, and superior longitudinal fasciculus 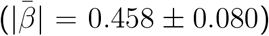 in ABCD (Appendices E-F,).

**Figure 7.**
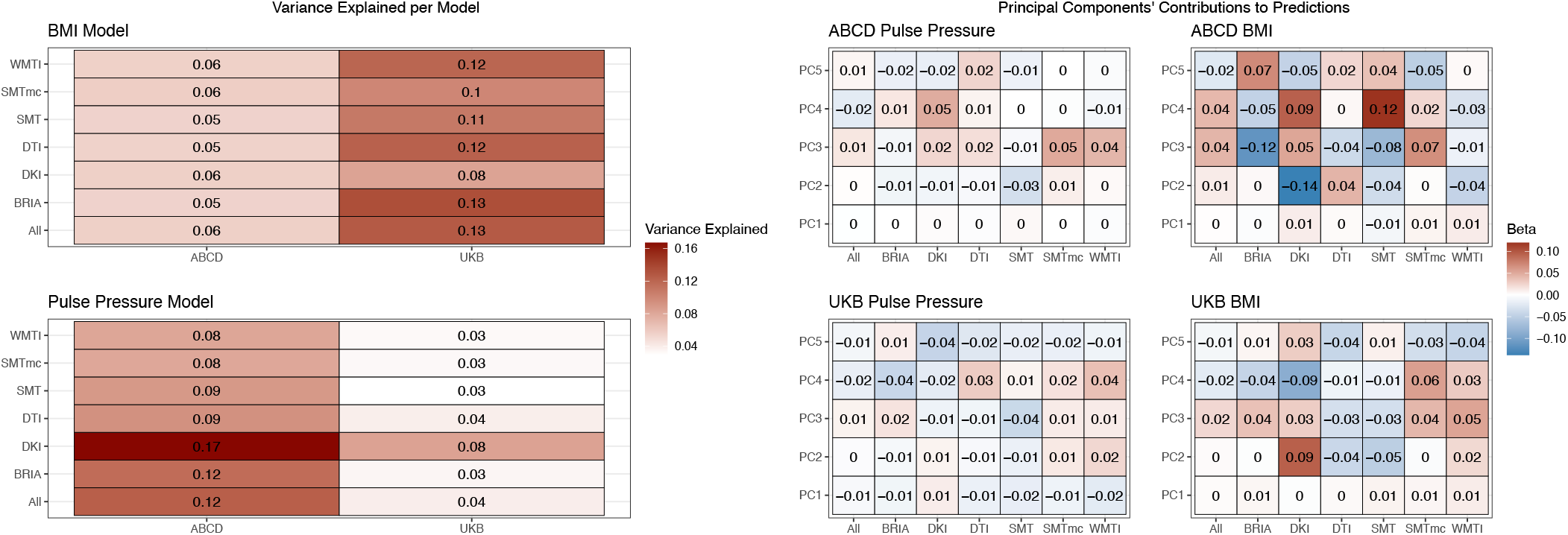
Variance in BMI and PP explained by principal components of the different diffusion approaches and the each component’s contribution to these predictions. BRIA = Bayesin Rotationally Invariant Approach. DKI = Diffusion Kurtosis Imaging. DTI = Diffusion Tensor Imaging. SMT = Spherical Mean Technique. SMTmc = SMT’s multi-compartment version. WMTI = White Matter Tract Integrity. Metrics for each of the approaches are specified in Appendix M. Standardized beta coefficients are presented.

Moreover, DKI was superior in predicting PP. Yet, none of the PCs were particularly strongly associated with BMI (Fig. 7).

### 3.5 Polygenic risk scores of psychiatric disorders and Alzheimer’s Disease

As a reference, we compared PGRS in the current samples (*N*_*ABCD*_ = 4, 906, *N*_*UKB*_ = 31, 408) to the rest of the ABCD dataset (*N*_*ABCD*_ = 6, 191) and UKB population (*N*_*UKB*_ = 377, 488), showing that PGRS were slightly lower in the present sample, but these differences were small (*d <* 0.07). PGRS with the largest difference were found for SCZ PGRS, being lower in the present imaging sample compared to the whole study population.

Effect sizes were small for the PGRS associations of mean skeleton metrics (|*β*| *<* 0.031), and even smaller for associations between tract-level PCs and PGRS (|*β*| *<* 0.024), with only few associations surviving multiple comparison corrections (Appendix N) and slightly stronger effects in the UKB compared to the ABCD data (Appendices O-P). Both single metrics and PCs of single diffusion approaches were similarly related to PGRS (Appendices O-P).

## 4 Discussion

In this paper, we presented an extensive analysis of the relationship of WM parameters from multiple dMRI approaches, at the tract- and skeleton-level, and a selection of clinically relevant metrics, including sex, age, brain asymmetry, and polygenic scores for common psychiatric disorders and Alzheimer’s disease. We examined one mid-life to late adulthood sample (UKB) and one adolescent sample (ABCD), with different biological characteristics. We described sample-specific sex differences in WM, WM asymmetries, and WM associations with pulse pressure, body-mass index, and polygenic risk scores. Our analyses highlight that multi-compartment dMRI approaches can add information over single-compartment approaches, such as DTI, when associating several geno- and phenotypes. For example, variability in microstructural asymmetries that are apparent in the ABCD (*d* = −3.4 to 3.7,) but not in the UKB *d* = 0 to −1.9) is particularly highlighted by the Bayesian multicompartment approach BRIA. Hence, stronger overall sex or age prediction accuracies across samples can be achieved, e.g. WMTI sex-prediction accuracy *>* 62%, BRIA age_*UKB*_ R^2^=0.36. At the same time, multi-compartment metrics add specificity and thereby biological meaning over mesostructure metrics, such as DTI fractional anisotropy or radial/axial/mean diffusivity.

### 4.1 Sex classifications and age-associations

Our analysis revealed dMRI approaches reflect sex differences fairly well, yet with some variations depending on the dataset. DKI and BRIA performed better in the ABCD dataset, while WMTI and DTI were superior in the UKB dataset. The variance in predictive accuracy between these datasets might be attributable to differences in participant demographics, age, or scanner-related effects. Determining a meaningful spatial pattern which describes sex-predictions was difficult and the highlighted tract contributions are to be interpreted with care (Appendices C-L).

Similarly, we examined variations in which metrics loaded on age-predicting components. That suggested that the strength of associations between WM metrics and different phenotypes might change across life, reflecting age-related changes in various phenoypes analogously. Nevertheless, we found that the cingulate tract was of overarching relevance for both age and sex predictions and other phenotype associations. Our findings also outline that all diffusion metrics follow individual age-associations in accordance with previous studies (Beck et al., 2021; Korbmacher, de Lange, et al., 2023; Korbmacher, van der Meer, Beck, Askeland-Gjerde, et al., 2024; Westlye et al., 2010). Sex differences in WM microstructure reported in the literature indicate varying effects in children and adolescents (Lawrence et al., 2023; López-Vicente et al., 2021) and adults (Eikenes et al., 2023; Korbmacher, de Lange, et al., 2023), and, for example, WM microstructure asymmetries were found to be similar between males and females (Korbmacher, van der Meer, Beck, de Lange, et al., 2024). However, ageing processes reflected by microstructure parametrisation differed significantly between sexes across conventional and advanced dMRI approaches (Korbmacher, van der Meer, Beck, Askeland-Gjerde, et al., 2024). The dMRI approaches which served best for sex classification in the current study (Fig. 2), contained also skeleton-level metrics showing the strongest sex differences in ageing in a previous study (Korbmacher, van der Meer, Beck, Askeland-Gjerde, et al., 2024). Similar to previously obtained results using different multi-compartment approaches (Lawrence et al., 2021), we conclude that advanced dMRI models might be more sensitive in detecting sex difference than conventional approaches, such as DTI.

BRIA explained most variance in WM age-associations, particularly in UKB (up to *R*^2^ =36%; Fig. 3). This model’s capacity to dissect intra- and extra-axonal compartments may provide more sensitive insights into age-related WM degeneration associated with axonal thinning and demyelination. For example, the cerebrospinal fluid fraction and microscopic diffusivity presented the strongest age-associations on the skeleton level (*r* = 0.49) in UKB, suggesting an age-related loss of neuronal tissue. In contrast, the observed age-associations in ABCD showcase WM differentiation during adolescence highlighted by lower diffusivity and extra-axonal water fractions, and higher anisotropy and intra-axonal water at a higher age, as previously presented in DTI (Westlye et al., 2010). These associations largely reverse in midlife to senescence (Fig. 4). Age-related changes in WM microstructure are well-documented, and advanced dMRI adds crucial information to simpler approaches like DTI (Beck et al., 2021; Korbmacher, de Lange, et al., 2023; Korbmacher, van der Meer, Beck, Askeland-Gjerde, et al., 2024). BRIA might be particularly useful to describe these processes due to the multiple different compartments. While the differences in diffusivity between intra- and extra-axonal space are highlighted across multi-compartment approaches, BRIA metrics might be able to provide additional information. For example, the positive age-associations of microscopic axial diffusivity in adolescence and then stronger positive association in senescence adds biologically plausible, yet additional detail to the distribution of diffusivity across compartments, likely reflecting both differentiation earlier in life and degeneration later in life (Fig. 4).

### 4.2 Tract asymmetry

There are a range of studies highlighting WM tract asymmetries (Arun et al., 2021; De Schotten et al., 2011). However, additional large-scale investigation is required to shed light on the mesostructural and microstructural WM asymmetry (Kong et al., 2020). One previous large-scale study in adults from mid-life to senescence demonstrated that asymmetry is present for most radiomic features, extracted from multimodal MRI, across brain regions (Korbmacher, van der Meer, Beck, de Lange, et al., 2024). To our knowledge, there are, however, no previous large-scale investigations of WM asymmetry in both children and adolescents. Here, we provide such overview by conducting a well-powered investigation of asymmetry separately for each of these age-groups and expressed as left-right differences. Asymmetry-derived findings might reflect important neurodevelopmental steps, for instance, the lateralisation of functions, such as attention, memory, and language (Barrick et al., 2007). Most of the asymmetry differences appeared more left-ward in the UKB/senescence sample, compared to the ABCD/adolescence sample (Fig. 5). BRIA metrics were most sensitive to these potential age differences in asymmetry. Including participants across the lifespan is strongly desired in order to provide additional biological meaning to the observed differences of WM asymmetry.

### 4.3 Cardiometabolic factors and polygenic risk score associations with WM microstructure

A growing body of evidence has presented associations between cardiometabolic factors and WM microstructure (Beck et al., 2022; Fuhrmann et al., 2019; Trofimova et al., 2023) and brain age derived from WM dMRI features (Beck et al., 2022; Korbmacher, Gurholt, et al., 2023). We associated body-derived parameters with multiple dMRI approaches’ WM parameters using UKB and ABCD data. We revealed stronger associations of several dMRI metrics with BMI than previously presented in the ABCD dataset (Li et al., 2023), and particularly highlight skeleton-level axial kurtosis associated with BMI. Tract-level analyses suggested that BMI was more strongly related with WM microstructure in the UKB sample than the ABCD for whole-skeleton averaged dMRI parameters. An opposite trend was observed for associations between PP and WM metrics, presenting weak associations with skeleton-level white matter metrics in the UKB. However, the associations highlighted by the principal components derived from tracts did not correspond with univariate associations between whole-skeleton averaged parameters, BMI and PP (compare Fig. and Fig. 7). Accordingly, neither a clear factor structure or corresponding spatial pattern based on loadings could be established. Yet, our findings suggest that body-derived measures could be considered as a proxy for different ageing manifestations, i.e. they might have a detectable relation with WM structure and associated changes in the brain tissue. These brain-body relationships offer unique potential to establish a better understanding of developmental and ageing processes. We revealed that imaging data represent sub-cohorts of the ABCD and UKB datasets which had significantly lower polygenic risk scores than the rest of the respective samples with the exception of ASD. Our findings reflect previously reported small associations between WM and PGRS in the UKB (Korbmacher, van der Meer, Beck, Askeland-Gjerde, et al., 2024) and the ABCD datasets (Fernandez-Cabello et al., 2022).

### 4.4 Limitations

The restricted age range in the ABCD dataset and the cross-sectional study design limit the ability to interpret the findings towards ageing trends. Moreover, throughout ageing, also the clinical significance of the examined markers might change. For example, high BMI and blood pressure might be more reliable markers of cardiometabolic disorders in older adults than in adolescents. Hence, when comparing the presented WM associations one needs to keep in mind the changes of both the examined WM and associated variables, which are likely non-linear. Additionally, during maturation and ageing, interaction effects between the observed variables cannot be excluded. Additional analyses of such effects are war-ranted. Lastely, the utilized acquisition protocols were similar to each other. Using different acquisition protocols, for example utilizing higher b-values, might allow to examine different effects which we were not able not observed here.

In conclusion, we analysed WM using multiple conventional and advanced dMRI approaches and related resulting parameters to metrics which are relevant to a broad range of research and clinical questions, including sex, age, brain asymmetry, BMI, PP, and polygenicity of psychiatric disorders and Alzheimer’s. Our findings demonstrate that the synergy of parameters derived from advanced dMRI approaches contributes to explanation of both WM organisation and its association with the included geno- and phenotypes. This highlights the utility of multi-compartment dMRI approaches, which contain more parameters than single compartment approaches, allowing for microstructure characterisations. Moreover, the congruence in the results for the used dMRI approaches in the two datasets increases the reproducibility of the findings and adds biological detail for explaining brain maturation processes.

## Data and Code Availability

Data are available from the UK Biobank https://www.ukbiobank.ac.uk and the ABCD Study https://abcdstudy.org after application. Code is available from GitHub https://github.com/MaxKorbmacher/ dMRI approach comparison.

## Author Contributions

M.K..: Study design, Software, Formal analysis, Visualizations, Project administration, Writing—original draft, Writing—review & editing. M.T.: Writing—review & editing. G.P.: Writing—review & editing. D.v.d.M.: Software, Writing – review & editing. M.-Y.W.: Writing—review & editing. O.A.A.: Writing—review & editing, Funding acquisition. L.T.W.: Writing—review & editing, Funding acquisition. I.I.M.: Study design, Data preprocessing and quality control, Writing—review & editing, Funding acquisition.

## Acknowledgements

We want to thank all UK Biobank and ABCD study facilitators and participants. Unfortunately, the most recent changes in UKB policy urge us to delete all UKB data from our servers limiting the computational reproducibility of our analyses and future research based on our dMRI data.

## Ethical Approval

This study was approved by the Norwegian Ethics Commission REK 567301, PVO 17/21624 (Ole Andreassen).

The study has been conducted using UKB data under Application 27412. UKB has received ethics approval from the National Health Service National Research Ethics Service (ref 11/NW/0382).

For the ABCD data, a centralised institutional review board approval of procedures was obtained from the University of California, San Diego. Written informed consent was obtained by a parent or guardian, and assent from the participants, before partaking in the ABCD study.

The work was performed on the Service for Sensitive Data (TSD) platform, owned by the University of Oslo, operated and developed by the TSD service group at the University of Oslo IT-Department (USIT). Computations were performed using resources provided by UNINETT Sigma2 – the National Infrastructure for High Performance Computing and Data Storage in Norway.

## Supplementary Material

Supplementary tables can be found in the GitHub repository https://github.com/MaxKorbmacher/dMRI_approach_comparison.

