## Supplement for "White matter microstructure links with brain, bodily and genetic attributes in adolescence, mid- and late life"

### Appendix

#### A Variance explained by each of the first 10 principal components of each diffusion approaches and their combination in the ABCD data.

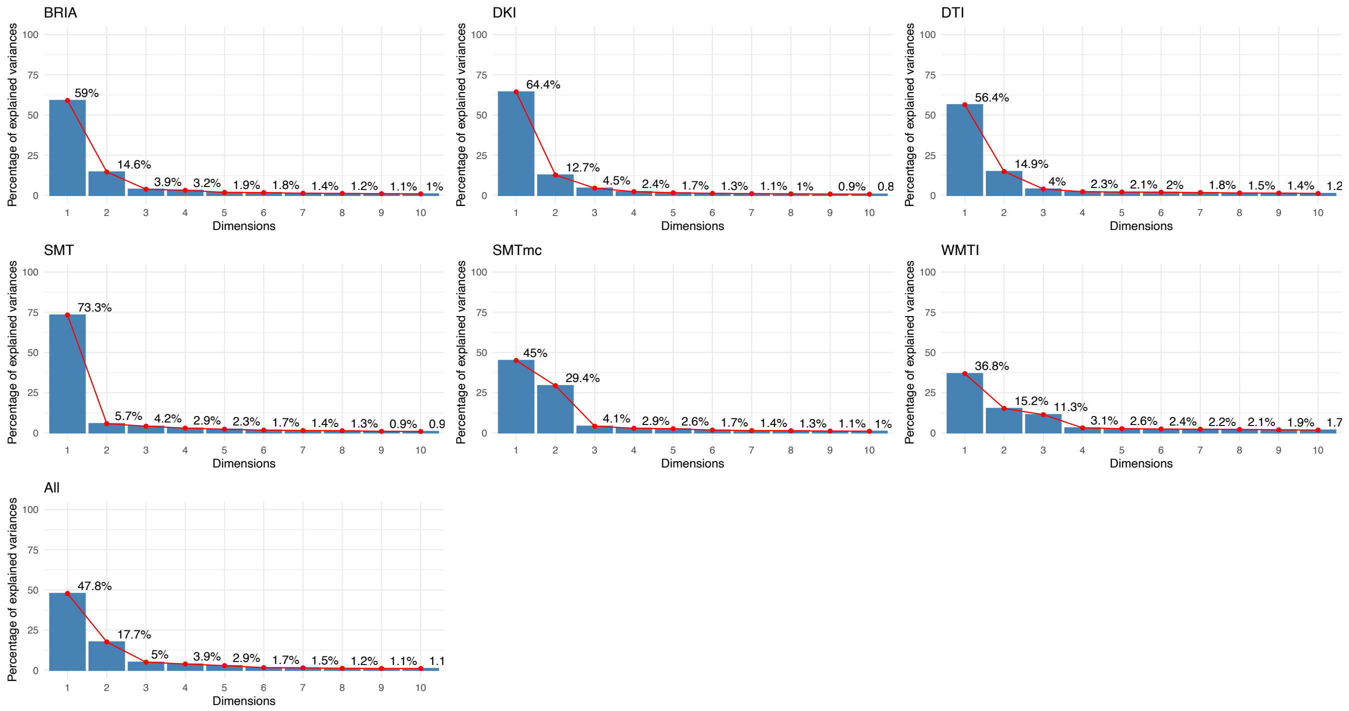

### B Variance explained by each of the first 10 principal components of each diffusion approaches and their combination in the UK Biobank data.

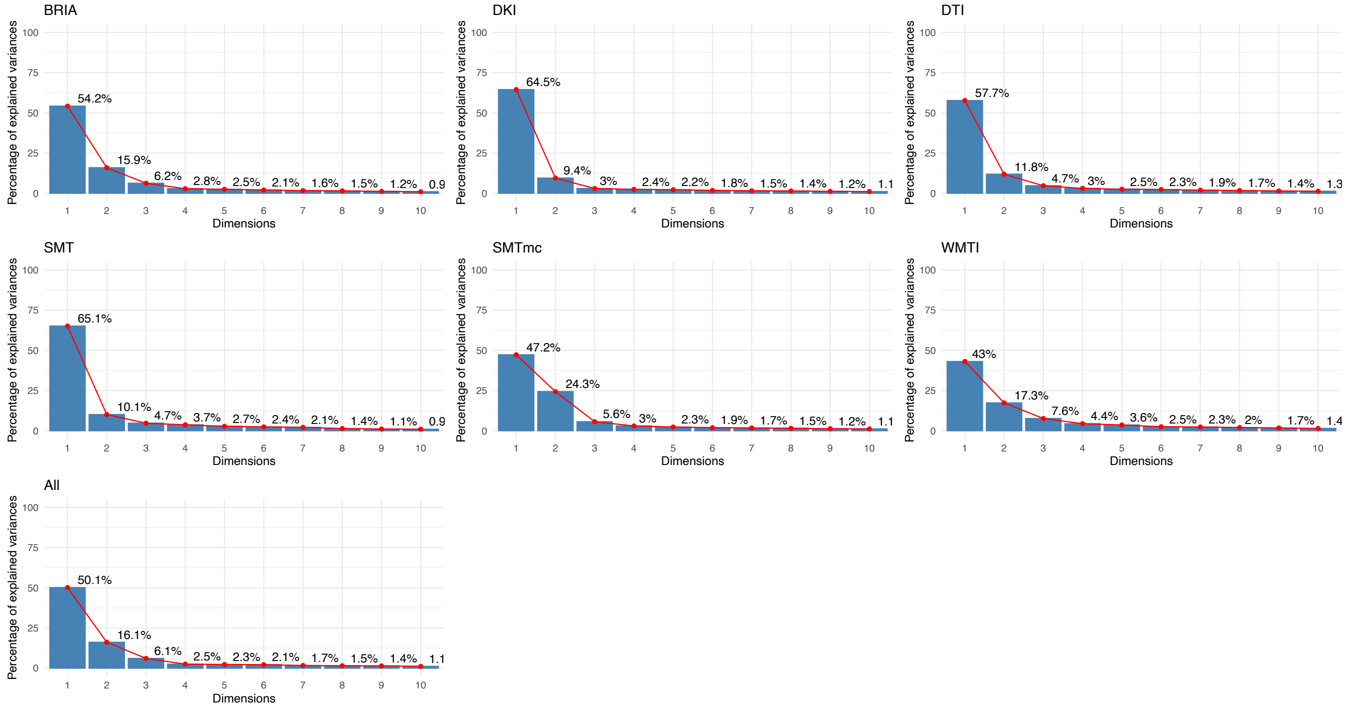

### C Corrected correlations between individual tract-level scalar estimates and their first principal component in ABCD data

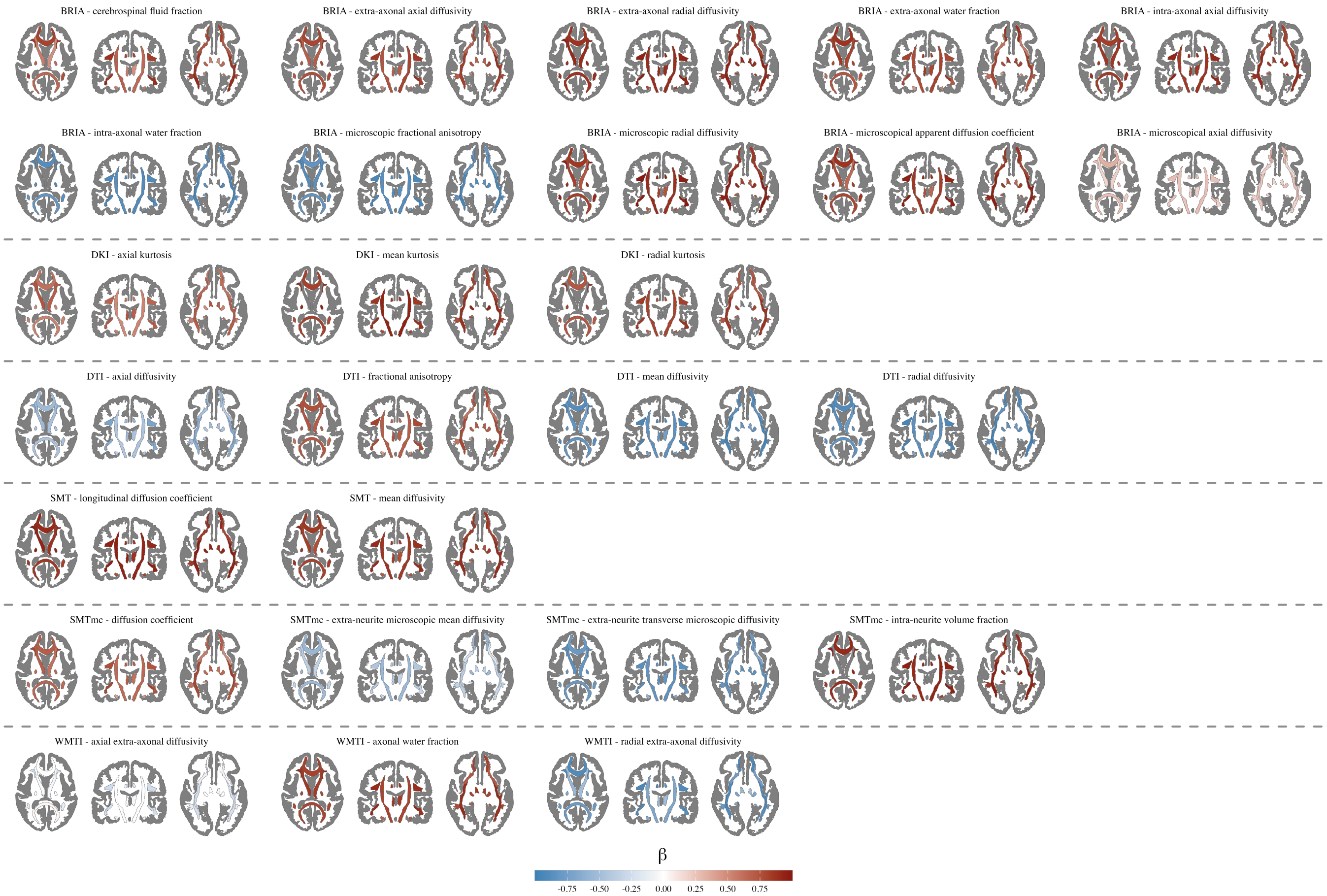

### D Corrected correlations between individual tract-level scalar estimates and their second principal component in ABCD data

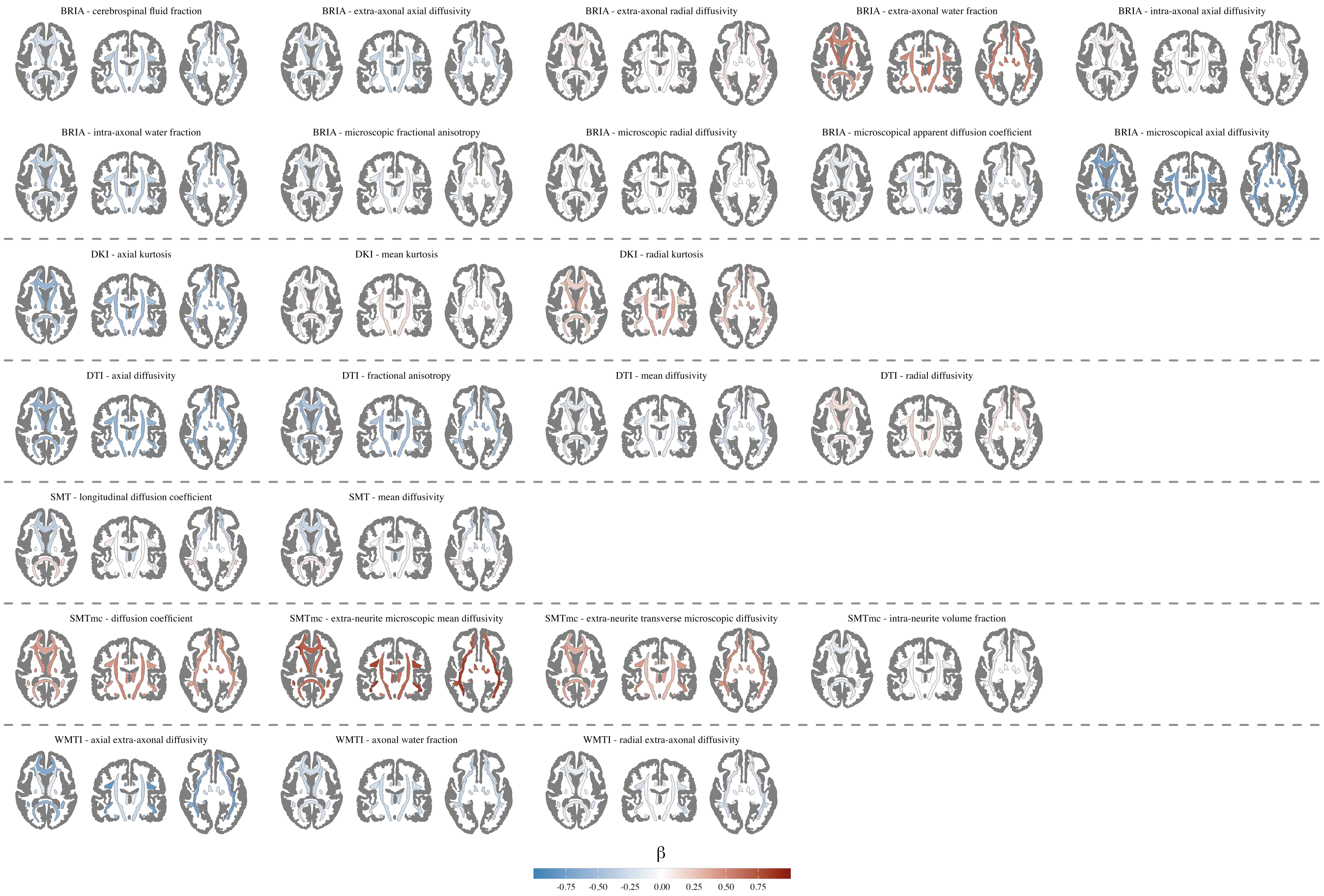

### E Corrected correlations between individual tract-level scalar estimates and their third principal component in ABCD data

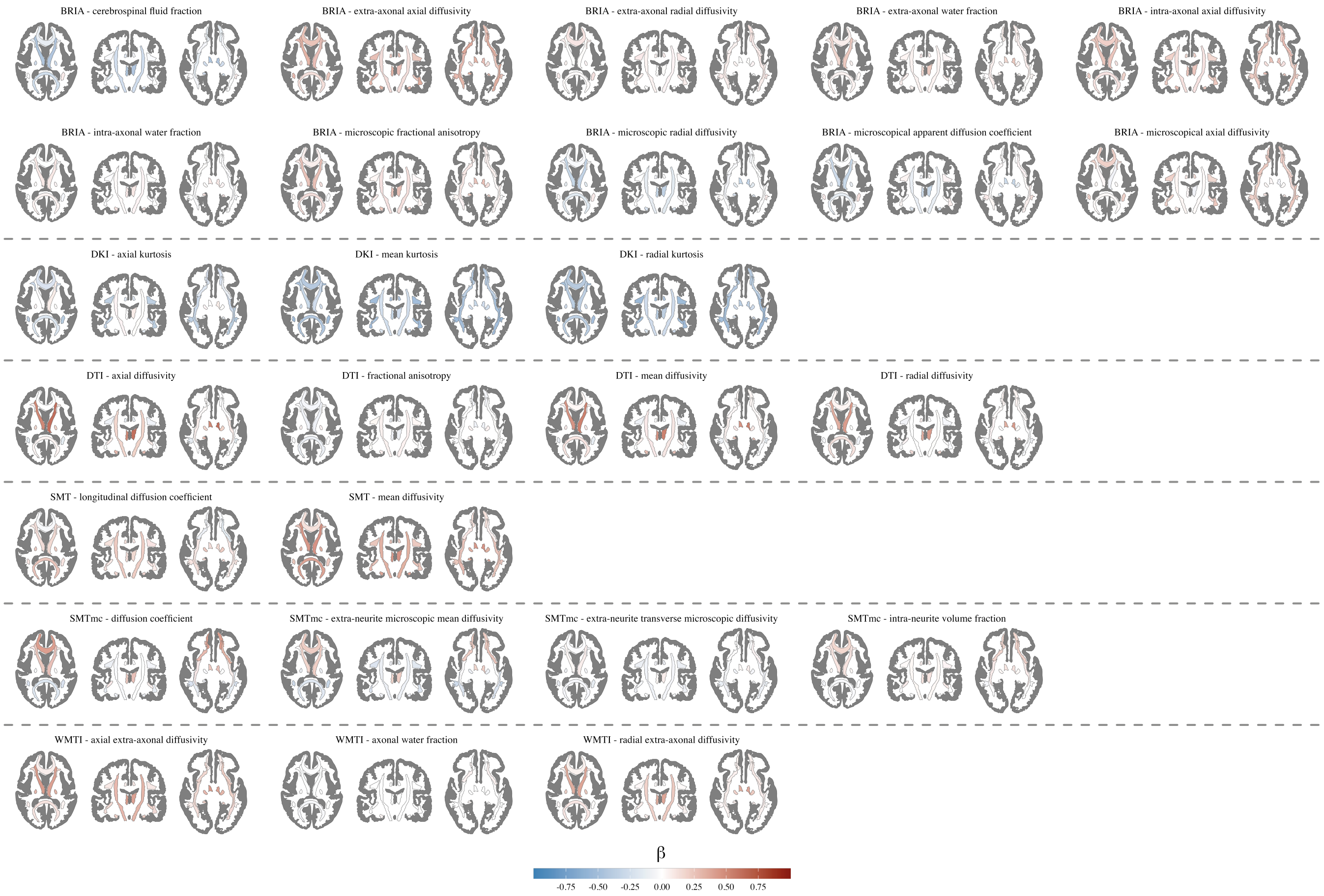

### F Corrected correlations between individual tract-level scalar estimates and their fourth principal component in ABCD data

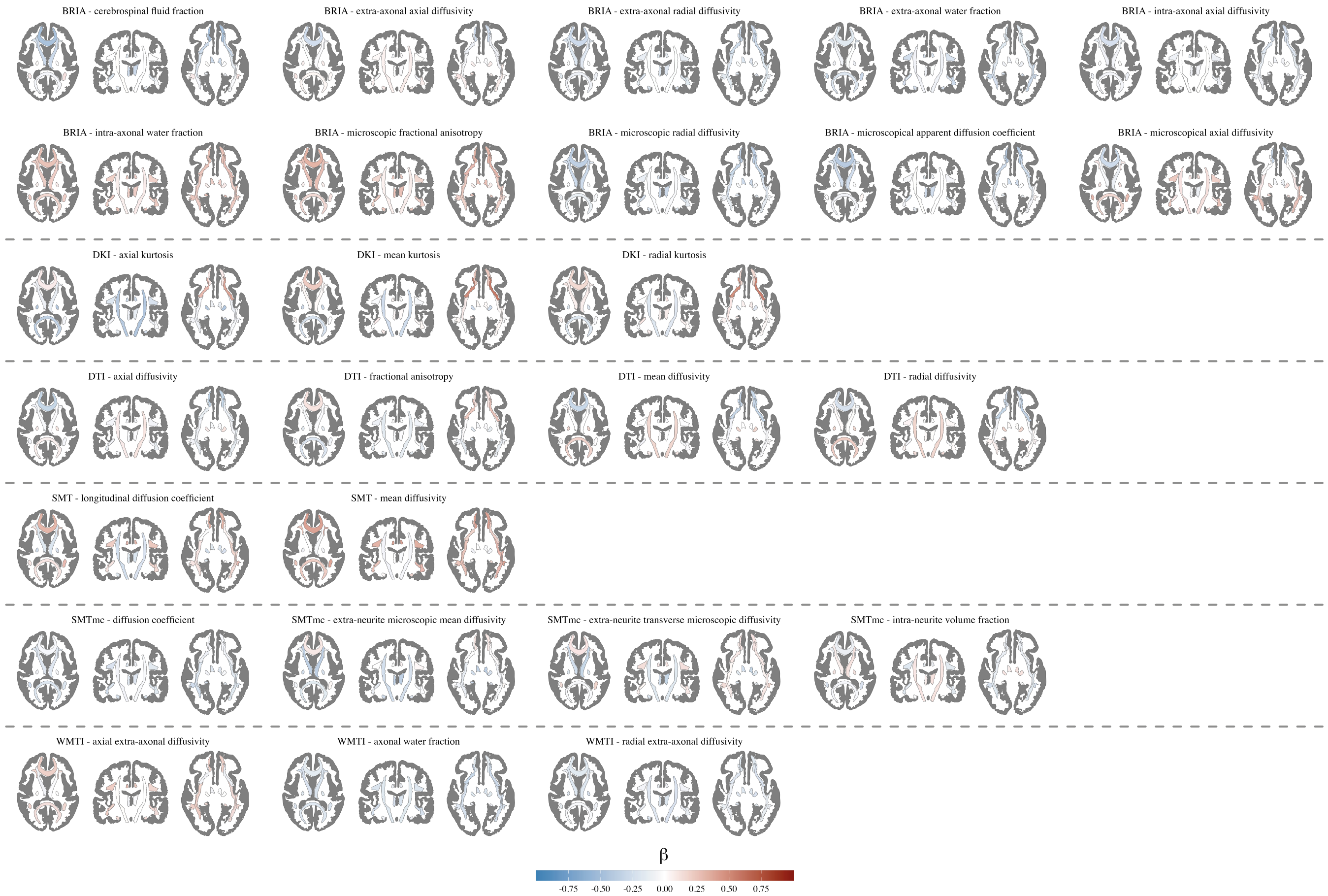

### G Corrected correlations between individual tract-level scalar estimates and their fifth principal component in ABCD data

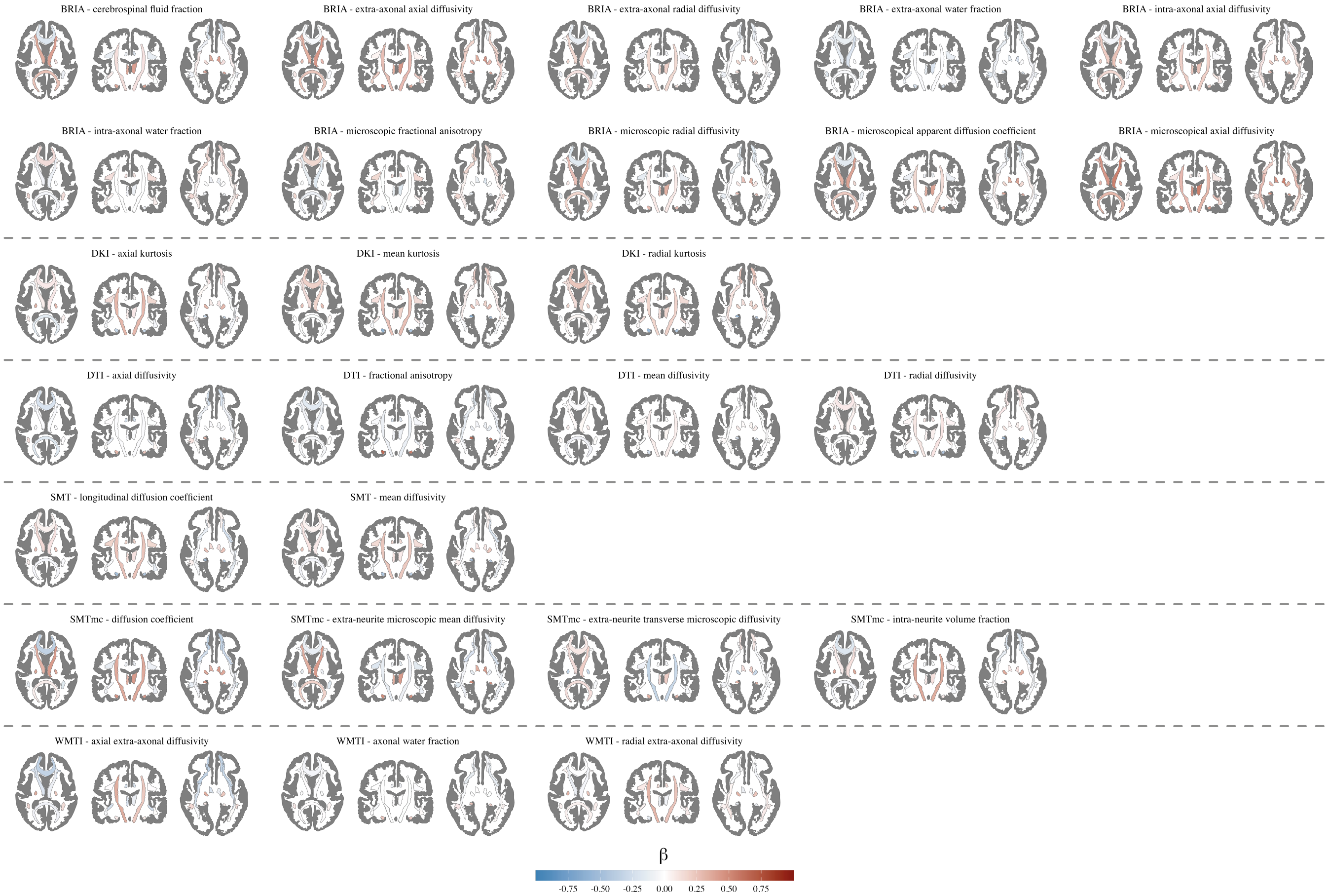

### H Corrected correlations between individual tract-level scalar estimates and their first principal component in UKB data

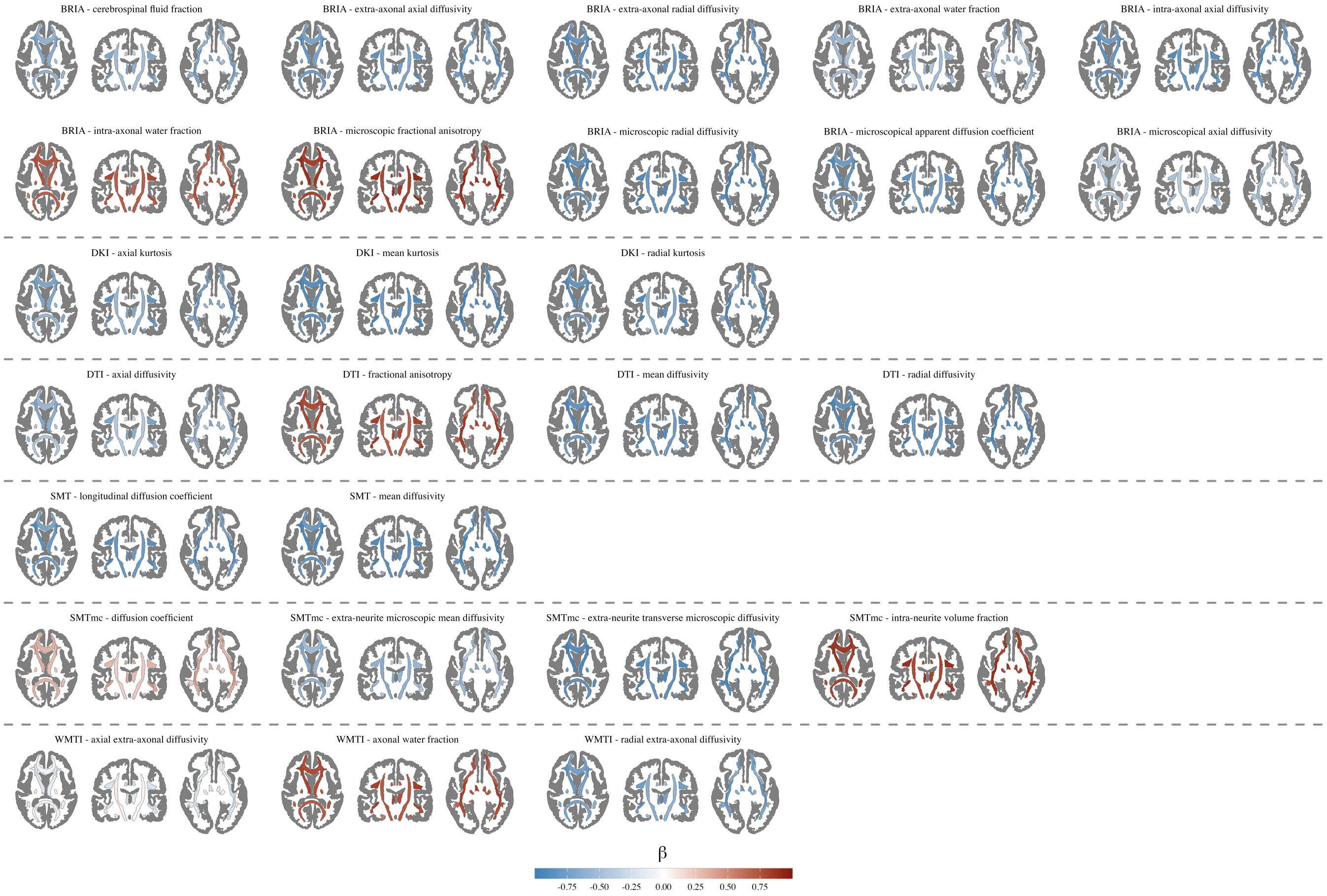

### I Corrected correlations between individual tract-level scalar estimates and their second principal component in UKB data

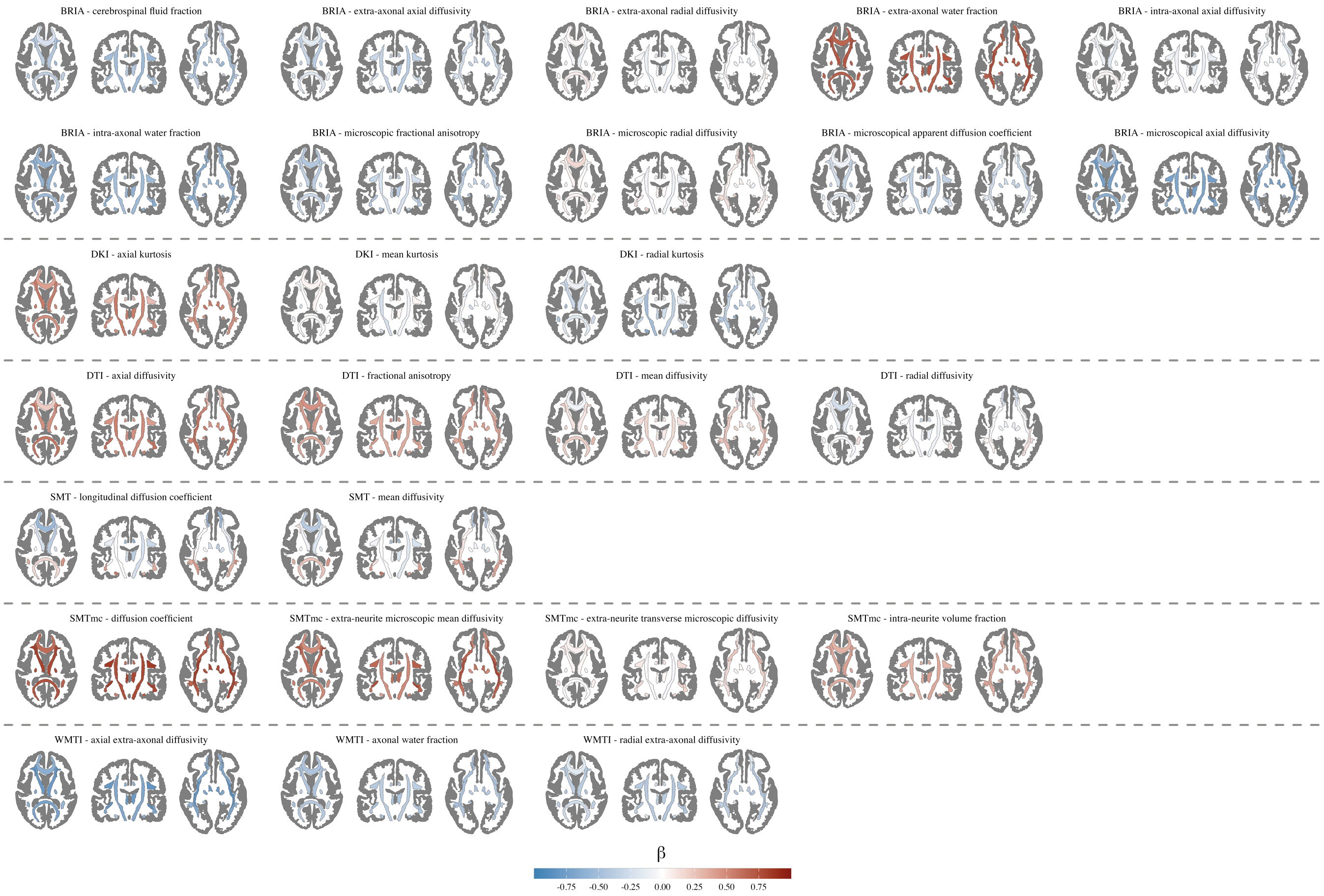

### J Corrected correlations between individual tract-level scalar estimates and their third principal component in UKB data

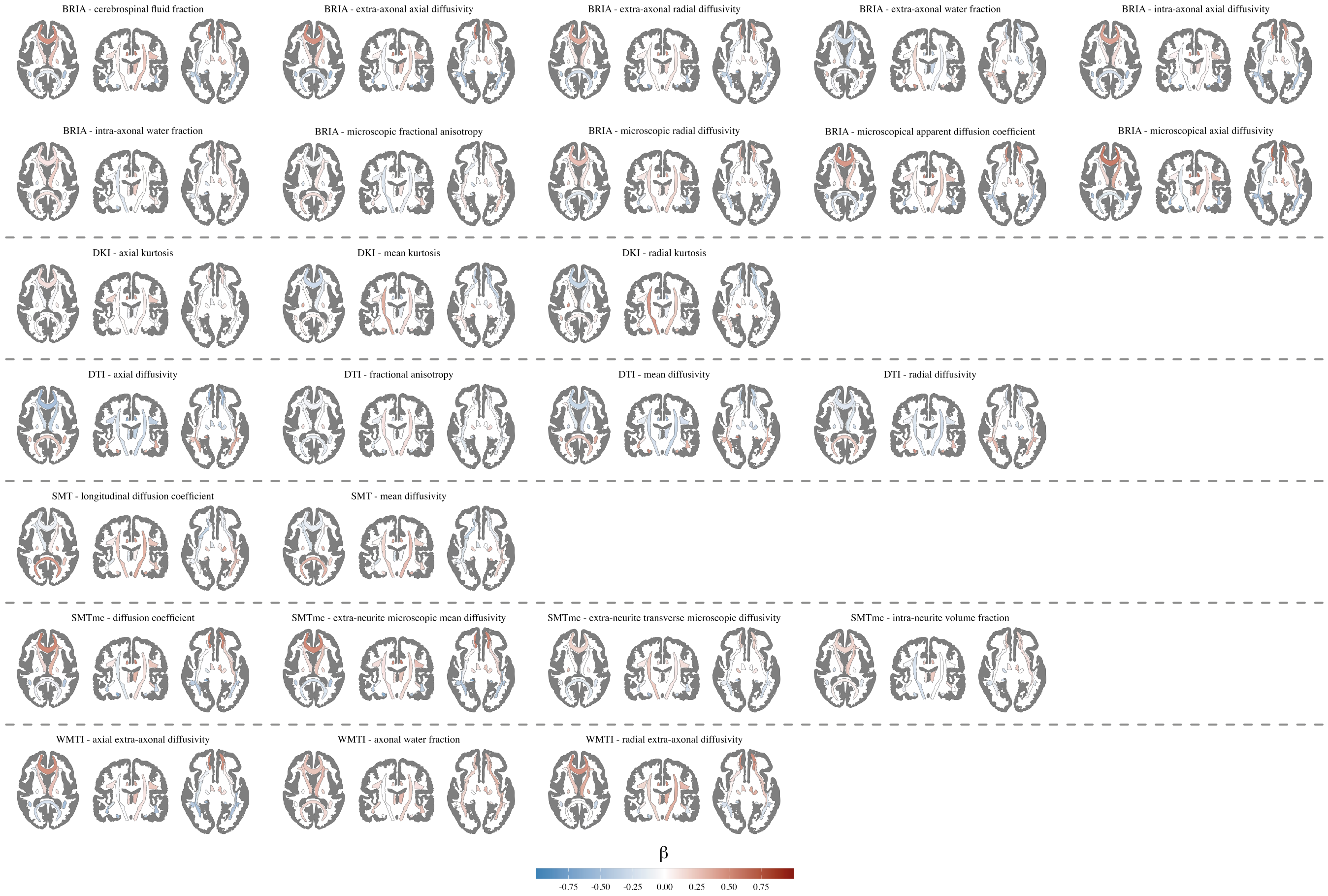

### K Corrected correlations between individual tract-level scalar estimates and their fourth principal component in UKB data

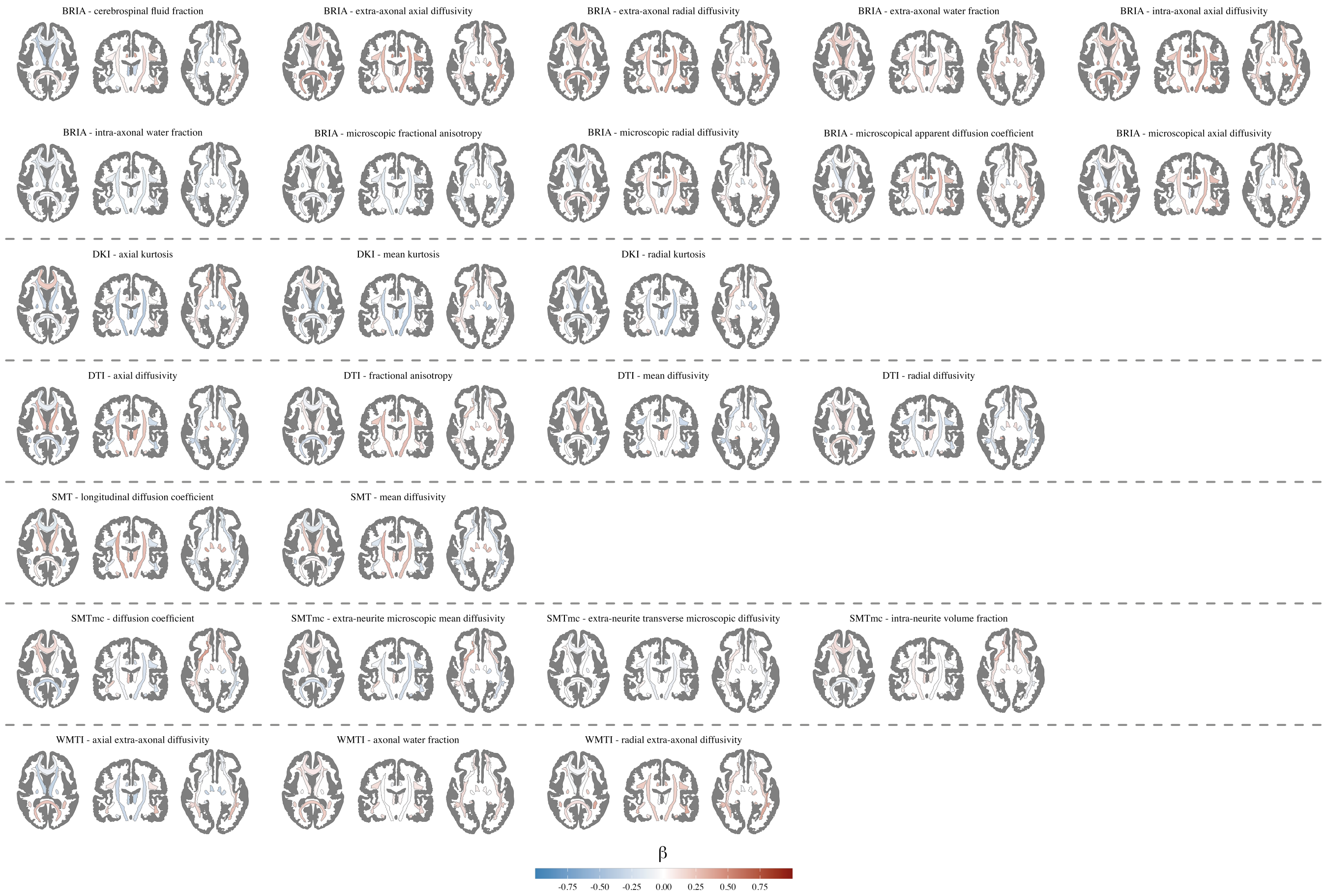

### L Corrected correlations between individual tract-level scalar estimates and their fifth principal component in UKB data

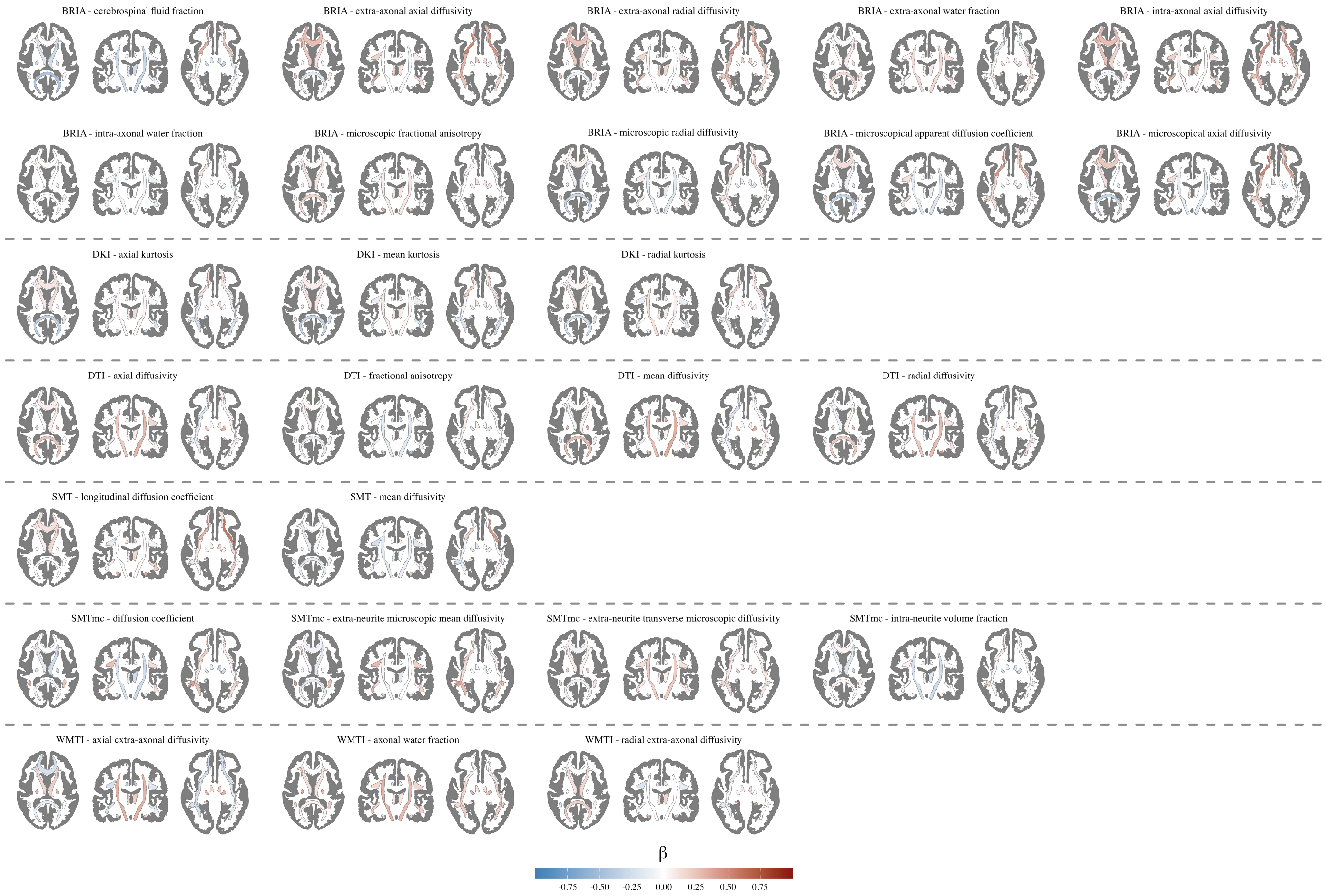

### M Overview of the used diffusion approaches and contained parameters.

| Diffusion Approach | parameter |
|---|---|
| Bayesian Rotationally Invariant Approach (BRIA) | |
| (Reisert et al., 2017) | intra-axonal axial diffusivity (DAX intra) |
| | extra-axonal radial diffusivity (DRAD extra) |
| | microscopic fractional anisotropy (micro FA) |
| | extra-axonal axial diffusivity (DAX extra) |
| | intra-axonal water fraction (V intra) |
| | extra-axonal water fraction (V extra) |
| | cerebrospinal fluid fraction (vCSF) |
| | microscopical axial diffusivity (micro AX) |
| | microscopic radial diffusivity (micro RD) |
| | microscopical apparent diffusion coefficient (micro ADC) |
| Diffusion Kurtosis Imaging | |
| (Fieremans et al., 2011; Jensen et al., 2005) | mean kurtosis (MK) |
| | radial kurtosis (RK) |
| | axial kurtosis (AK) |
| Diffusion Tensor Imaging (DTI) | |
| (Basser et al., 1994) | fractional anisotropy (FA) |
| | axial diffusivity (AD) |
| | mean diffusivity (MD) |
| | radial diffusivity (RD) |
| Spherical Mean Technique (SMT) | |
| (Kaden, Kruggel, & Alexander, 2016) | fractional anisotropy (SMT FA) [Not included] |
| | mean diffusivity (SMT md) |
| | transverse diffusion coefficient (SMT trans) [Not included] |
| | longitudinal diffusion coefficient (SMT long) |
| Multi-compartment Spherical Mean Technique (SMTmc) | |
| (Kaden, Kelm, et al., 2016) | extra-neurite microscopic mean diffusivity (SMTmc extra md) |
| | extra-neurite transverse microscopic diffusivity (SMTmc extra trans) |
| | mc SMTdiffusion coefficient (SMT mcd) |
| | intra-neurite volume fraction (SMTmc intra) |
| White Matter Tract Integrity | |
| (WMTI) (Fieremans et al., 2011) | axonal water fraction (AWF) |
| | radial extra-axonal diffusivity (radEAD) |
| | axial extra-axonal diffusivity (axEAD) |

### N Associations between PGRS and mean WM skeleton metrics as well as PCs which survive the Bonferroni correction procedure

Only the PCs of DKI were significantly related to BIP PGRS in ABCD ($\beta = -0.009, p_{corrected} = 0.035$) and SCZ in UKB ($\beta = -0.023, p_{corrected} = 0.042$), as well as the SMT PC with MDD in UKB ($\beta = -0.006, p_{corrected} = 0.046$). These associations were not as well powered (at 95%) as aimed for in the Methods section and hence not reported in the main text. The associations between mean skeleton values and PGRS can be found in the table below.

| Group | Variable | Uncorrected p | Corrected p |
|---|---|---|---|
| UKB-SCZ | BRIA-Vintra | 2.592794e-05 | 0.0183569842 |
| ABCD-BIP | BRIA-vextra | 2.562437e-05 | 0.0181420547 |
| UKB-BIP | BRIA-microRD | 4.539597e-05 | 0.0321403465 |
| UKB-BIP | BRIA-microFA | 2.389440e-05 | 0.0169172368 |
| UKB-SCZ | BRIA-microFA | 1.357681e-05 | 0.0096123823 |
| UKB-SCZ | DKI-MK | 8.522015e-06 | 0.0060335865 |
| UKB-SCZ | DKI-RK | 3.321437e-07 | 0.0002351577 |
| ABCD-BIP | DKI-AK | 9.426733e-06 | 0.0066741269 |
| UKB-SCZ | DTI-FA | 6.056883e-06 | 0.0042882735 |
| UKB-MDD | DTI-RD | 6.286053e-05 | 0.0445052532 |
| UKB-MDD | SMT-MD | 3.216320e-05 | 0.0227715461 |
| UKB-SCZ | SMTmc-intra | 3.667747e-05 | 0.0259676501 |
| ABCD-BIP | SMTmc-intra | 4.360449e-05 | 0.0308719810 |
| UKB-SCZ | WMTI-AWF | 1.734435e-05 | 0.0122797989 |

### O Associations between PGRS and mean WM skeleton metrics

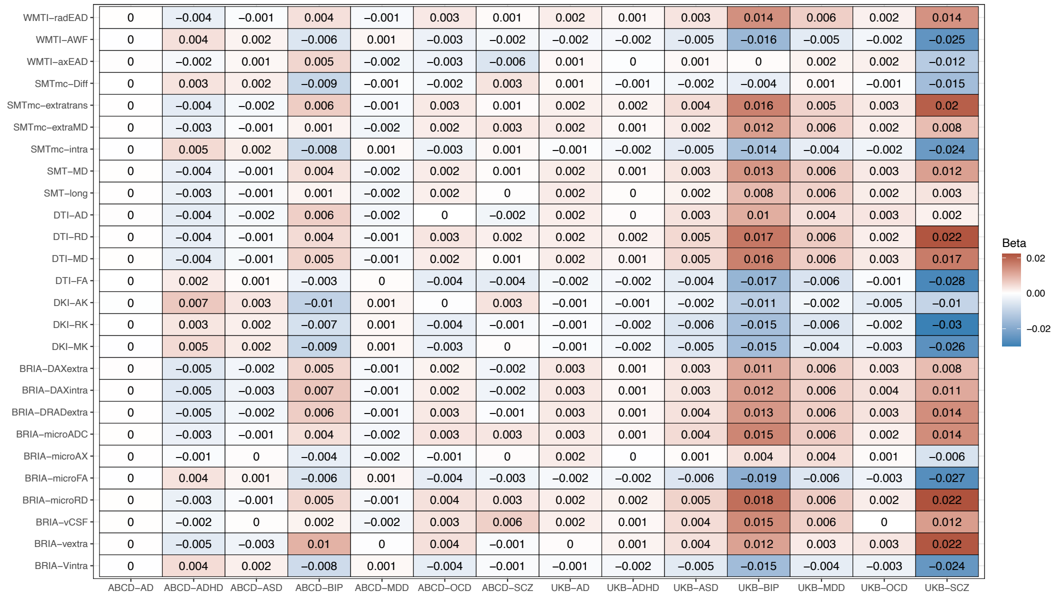


Figure 8: The polygenic risk scores (PGRS) of psychiatric disorders, the PGRS of Alzheimer's Disease and skeleton-level metrics. BRIA = Bayesin Rotationally Invariant Approach. DKI = Diffusion Kurtosis Imaging. DTI = Diffusion Tensor Imaging. SMT = Spherical Mean Technique. SMTmc = SMT's multi-compartment version. WMTI = White Matter Tract Integrity. Metrics for each of the approaches are specified in Appendix M.

### P Associations between PGRS and mean Principal Components of WM tract metrics

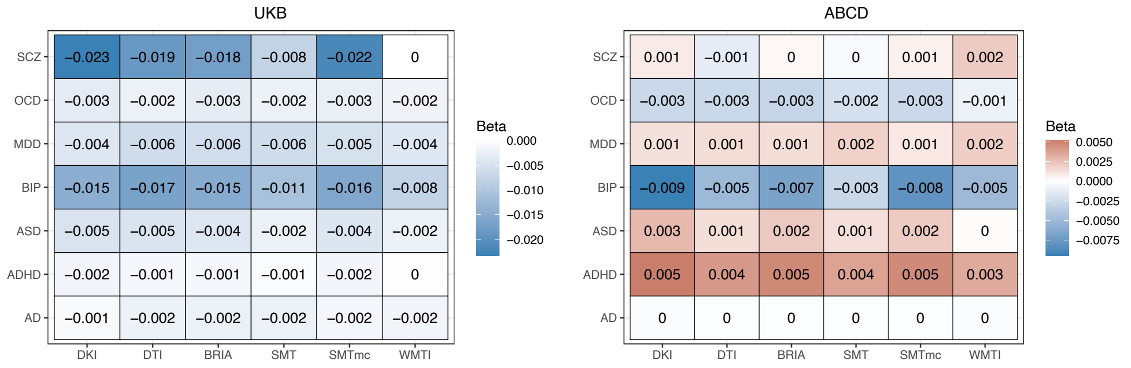


Figure 9: Associations between polygenic risk scores (PGRS) of psychiatric disorders, the PGRS of Alzheimer's Disease and tract-level metrics' principal components. BRIA = Bayesin Rotationally Invariant Approach. DKI = Diffusion Kurtosis Imaging. DTI = Diffusion Tensor Imaging. SMT = Spherical Mean Technique. SMTmc = SMT's multi-compartment version. WMTI = White Matter Tract Integrity. Metrics for each of the approaches are specified in Appendix M.
